# A low repeated dose of Δ9-tetrahydrocannabinol affects memory performance through serotonergic signalling in mice

**DOI:** 10.1101/2021.05.31.446448

**Authors:** Lorena Galera-López, Victòria Salgado-Mendialdúa, Estefanía Moreno, Araceli Bergadà-Martínez, Alexander F. Hoffman, Irene Manzanares-Sierra, Arnau Busquets-Garcia, Vicent Casadó, Carl R. Lupica, Rafael Maldonado, Andrés Ozaita

## Abstract

Cannabis is the most widely used illicit drug worldwide. Its principal psychoactive component, Δ9-tetrahydrocannabinol (THC), acts as a partial agonist of the main cannabinoid receptor in the brain, the cannabinoid type-1 receptor (CB1R), that is responsible for the central effects of THC including memory impairment. CB1Rs may form heterodimers with the serotonin 5-HT2A receptor (5-HT2AR) which were found responsible for the memory impairment produced by acute high dose of THC in mice. In this study we investigated whether a repeated low dose of THC (1 mg/kg), with no acute consequence on memory performance, could eventually have deleterious cognitive effects. We found that this dose of THC impaired novel object-recognition memory and fear conditioning memory 24 h after the last of 7 consecutive daily treatments. At that time, a general enhancement of c-Fos expression was also observed in several brain regions of THC-exposed animals, as well as a decreased dendritic spine density on hippocampal CA1 pyramidal neurons that was accompanied by reduced long-term potentiation (LTP) at Schaffer collateral-CA1 synapses. Interestingly, an up-regulation in the expression of CB1R/5-HT2AR heterodimers was observed in the hippocampus after THC exposure and pre-treatment with the 5-HT2AR antagonist MDL 100,907 (0.01 mg/kg) prevented the enhanced heterodimerization and the THC-associated memory impairment. Together, these results reveal the significance of serotonergic signalling through 5-HT2ARs in the memory-impairing effects of repeated low doses of THC.

## Introduction

Preparations from the Cannabis sativa plant have been consumed for medical and recreational purposes over thousands of years (Bonini et al., 2018), and cannabis remains the most used illicit drug worldwide (Zarei et al., 2020). From the more than 400 compounds, including over 120 phytocannabinoids, that have been extracted from cannabis, Δ9-tethahydrocannabinol (THC) has been identified as its main psychoactive component (dos Santos et al., 2021). THC has been extensively studied due to its potential therapeutic applications including analgesia properties (King et al., 2017; Lötsch et al., 2018; Escudero-Lara et al., 2020), anxiolytic-like effects, and neuroprotective effects (Fraguas-Sánchez and Torres-Suárez, 2018; Maroon and Bost, 2018), among others. However, THC also induces several undesirable effects such as anxiety (Kasten et al., 2017), dependence (Moreira and Lutz, 2008; Maldonado et al., 2011), and memory impairment (Puighermanal et al., 2012; Volkow et al., 2016). Central effects of THC, including memory impairment, are mostly mediated by the cannabinoid type-1 receptor (CB1R), the main receptor of the endocannabinoid system, an important brain homeostatic system (Kano et al., 2009). Importantly, THC-induced impairments in object-recognition memory are completely absent in mice lacking CB1R (Puighermanal et al., 2009). The acute amnesic-like effects of THC in mice are dose dependent, and low doses of THC do not produce memory impairments in object recognition memory following an acute administration (Puighermanal et al., 2009). THC can also impair synaptic plasticity at a functional and structural level. Thus, 1 day after repeated treatment (7 d) with a relatively high dose of THC (10 mg/kg) CA1 dendritic spine density is decreased in the hippocampus and LTP is prevented (Hoffman et al., 2007; Chen et al., 2013). However, LTP is normal 1 day after a single THC treatment at the same dose (Hoffman et al., 2007). Although these studies highlight the potential for extended exposure to relatively high doses of THC to cause changes in hippocampal structure and function, presently it is unknown whether repeated exposure to low doses of THC, that might better represent those encountered by humans using cannabis, can affect memory and plasticity.

The CB1R is abundantly expressed in the central nervous system and controls the release of several major neurotransmitters and neuromodulators including γ-aminobutyric acid (GABA), glutamate, acetylcholine, noradrenaline, dopamine, cholecystokinin and serotonin (Lutz, 2020). Among those, a crosstalk between the endocannabinoid system and the serotonergic system has been described. For example, the activation of serotonin 5-HT2A receptors (5-HT2AR) induces the release of endocannabinoids, the endogenous ligands for cannabinoid receptors (Parrish and Nichols, 2006; Best and Regehr, 2008), and the serotonergic system has been proposed to be involved in several THC-induced effects including hypothermia (Malone and Taylor, 2001), catalepsy (Egashira et al., 2007), anxiety, sociability and memory (Viñals et al., 2015). Moreover, the memory disrupting effects of an acute high dose of THC in object recognition is prevented in mice lacking 5-HT2ARs, or those pre-treated with the 5-HT2AR antagonist MDL 100,907 (MDL) (Viñals et al., 2015). The interactions between serotonin and cannabinoid receptors may be reciprocal since mice lacking CB1Rs showed a reduced number of head twitches caused by the 5-HT2R agonist, 2,5-Dimethoxy-4-iodoamphetamine (DOI) (Mato et al., 2007) as well as a dysregulation in serotonergic activity in the prefrontal cortex (Aso et al., 2009).In further support of this interaction, heterodimers of CB1R and 5-HT2AR have been described in both mice and humans (Viñals et al., 2015; Galindo et al., 2018). Enhanced expression of CB1R/5-HT2AR heterodimers has also been found in human cannabis users (Galindo et al., 2018), and enhanced levels and altered signalling of CB1R/5-HT2AR heterodimers was observed in olfactory neuroepithelium cells of schizophrenia patients (Guinart et al., 2020). In mice, the presence of CB1R/5-HT2AR heterodimer has been described in CA3 region of the hippocampus, the cortex, and the striatum. Importantly, CB1R/5-HT2AR heterodimers appear to modulate specific acute effects of THC since an interference peptide that prevents heterodimer formation limited the memory impairment and anxiolytic effects of THC, whereas other classical effects of the phytocannabinoid were unaffected (Viñals et al., 2015).

In this study, we investigated the effect of repeated exposure to an acutely non-amnesic low-dose of THC (1 mg/kg) on cognitive function in mice and the possible involvement of 5-HT2ARs on those effects. We found that repeated low-dose THC resulted in specific deficits in non-emotional and emotional hippocampal long-term memory, as well as physiological and molecular alterations in hippocampal brain regions related to these cognitive deficits. Thus, a decrease in CA1 pyramidal neuron dendritic spine density and LTP at CA3-CA1 synapses were observed, as well as a general enhancement of c-Fos expression compared to control. We also report that CB1R/5-HT2AR heterodimer expression was increased in the CA3 region of the hippocampus following chronic low-dose THC, that antagonism of 5HT2ARs prevented the effect of THC on memory performance, and that MDL treatment before THC prevented the changes caused by THC alone. Altogether, our results indicate that long-term exposure to an acutely sub-effective dose of THC can produce memory impairment and cellular alterations in mice, and that these effects are mediated by 5-HT2AR.

## Materials and Methods

### Ethics

All animal procedures were conducted following ARRIVE (Animals in Research: Reporting In Vivo Experiments) guidelines (Kilkenny et al., 2010) and standard ethical guidelines (European Union Directive 2010/63/EU) and approved by the local ethical committee (Comitè Ètic d’Experimentació Animal-Parc de Recerca Biomèdica de Barcelona, CEEA-PRBB). All electrophysiological studies were approved by the NIDA IRP Animal Care and Use Committee and carried out in accordance with NIH guidelines.

### Animals

Young adult (8-12 weeks old) male Swiss albino (CD-1) mice (Charles River) and C57BL/6J (BL-6) mice (Charles River) were used for pharmacological approaches on behavioural, biochemical, and electrophysiological approaches. The transgenic line Tg(Thy1-EGFP) MJrs/J (Thy1-EGFP mice) in C57BL/6J background (Stock # 007788, The Jackson Laboratories) was used to study the density and morphology of dendritic spines.

Mice were housed in controlled environmental conditions (21 ± 1 °C temperature and 55 ± 10% humidity) and food and water were available ad libitum. All the experiments were performed during the light phase of a 12 h light/dark cycle (light on at 8 am; light off at 8 pm). Mice were habituated to the experimental room and handled for 1 week before starting the experiments. All behavioural experiments were conducted by an observer blind to the experimental conditions.

### Drug treatment

Delta9-tetrahydrocannabinol (THC) (THC-Pharm-GmbH) was diluted in 5 % ethanol, 5 % Cremophor-EL and 90 % saline and injected intraperitoneally (i.p.) at the dose of 1 mg/kg in a volume of 10 mL/kg of body weight. For electrophysiological studies, THC was provided by NIDA Drug Supply (Rockville). MDL 100,907 (MDL) (Sigma-Aldrich) was diluted in 0.26 % dimethyl sulfoxide (DMSO; Scharlau Chemie), 4.74 % ethanol, 5 % Cremophor-EL and 90 % saline at the dose of 0.01 mg/kg in a volume of 10 mL/kg of body weight 20 min before Veh/THC administration. The length of the sub-chronic administrations consisted of 6 d or 7 d of treatment, as indicated in the results section.

### Behavioural tests

#### Novel object recognition test

The novel object recognition (NOR) memory test is a behavioural task used to measure non-emotional declarative memory in mice. NOR test was performed as described before (Gomis-González et al., 2021) in a V-shaped maze (V-maze). On day 1, mice were habituated to the empty V-maze for 9 min (habituation phase). On day 2, two identical objects (familiar objects) were presented at the end of each corridor of the V-maze and mice were left to explore for 9 min before they were returned to their home cage (familiarization phase). On day 3 (24 h after the familiarization phase), mice were placed back in the V-maze where one of the familiar objects was replaced by a new object (novel object) to assess memory performance (test phase). The time spent exploring each of the objects (familiar and novel) during the test session was computed to calculate a discrimination index (DI = (Time Novel Object - Time Familiar Object)/(Time Novel Object + Time Familiar Object)), defining exploration as the orientation of the nose toward the object at a distance closer than 1 cm. A higher discrimination index is considered to reflect greater memory retention for the familiar object. Mice that explored <10 sec both objects during the test session or <2 sec one of the objects were excluded from the analysis. For this task, drug treatment was administered after the familiarization phase in acute and sub-chronic treatments. For the measurement of the effects after treatment withdrawal the treatment finished 24 h prior to the familiarization phase. For NOR task CD-1 mice were used (Suppl. Fig. 1a).

#### Consecutive NOR

The repeated assessment of NOR memory was carried out as previously described (Puighermanal et al., 2013). Briefly, the first memory assessment (T_1) on the consecutive NOR procedure was performed 24 h after the first administration of THC and just before the second drug administration. Memory was tested again 24 h later using the novel object explored the day before (now the familiar object) and a brand-new object (now the novel object, to be the familiar object the day after). In each test session, a discrimination index (DI) was calculated as previously explained. Mice received a total of 6 administrations for this specific task and a total of 10 NOR memory assessments were performed. For consecutive NOR task CD-1 mice were used.

#### Trace and Context fear conditioning

Fear conditioning (FC) is a behavioural task used to measure emotional memory in mice. Trace FC is composed of 2 phases. On day 1, mice were placed in a conditioning chamber with an electrifiable floor. After 2 min of free exploration mice were exposed to an auditory stimulus (conditioned stimulus, CS) for 1 min. Then, 15 sec after the end of the auditory stimulus mice received one foot shock (unconditioned stimulus, US: 2 sec, 0.35 mA intensity) (training phase). On day 2 (Trace FC test phase), 24 h after the training phase, mice were placed in a new environment (transparent cylinder). After 2 min of free exploration, the exact same sound that was presented during training (CS) was played for 1 min; during this time, the freezing behaviour was manually counted. 2 min after the sound end, mice were removed from the cylinder. The time mice showed freezing behaviour during exposure to CS was expressed as a percentage and interpreted as a memory strength index.

In some cases, Context FC was also measured. For this purpose, an additional phase was added to the previous protocol. On day 3 (Context FC test phase), 24 h after the Trace FC test phase, mice were placed for 5 min in absence of the shock in the same conditioning chamber used in the training phase on day 1 and the freezing behaviour was manually recorded.

For this task, drug treatment was administered after the training phase for acute and sub-chronic treatment. For the measurement of the effects after treatment withdrawal the treatment finished 24 h prior to the training phase. For FC tasks BL-6 animals were used (Suppl. Fig. 1a).

#### Spontaneous alternation task

Spontaneous alternation task is a behavioural test to measure spatial working memory. The maze is composed by three identical arms that intersected at 120° (Y-shaped; 6.5-cm width × 30-cm length × 15-cm height, Y-maze). Animals freely explore the Y-maze for 5 min, starting from the end of the same arm in the Y-maze facing the wall. Entries into all arms of the Y-maze were counted (traversing the head and two front paws was considered a valid entry) and the percentage of correct spontaneous alternations was calculated considering the sequential entries in all three arms divided by the total number of possible alternations (triplets) calculated as the total number of entries − 2. For this task, drug treatment was administered 24 h before both the acute and sub-chronic treatments. For the spontaneous alternation task CD-1 animals were used (Suppl. Fig. 1a).

#### Anxiety-like behaviour (0-maze and elevated plus maze)

Anxiety-like behaviour associated to the acute and chronic administration of THC was measured using the 0-maze and elevated plus maze. The 0-maze consisted of a black Plexiglas round shape corridor (5 cm wide), where 2 quarters of the maze were closed by walls (20 cm-high) and the other 2 quarters were open. The whole maze is elevated 30 cm above the floor. During 5 min mice freely explored the maze and the percentage of time spent in the open parts of the corridor were counted as a measure of anxiety-like behaviour. Animals that exit the maze during exploration were excluded. The elevated plus maze test consisted of a black Plexiglas apparatus with 4 arms (29 cm long x 5 cm wide), 2 closed arms with walls (20 cm high) and 2 open arms, set in cross from a neutral central square (5 x 5 cm) elevated 40 cm above the floor. Light intensity in the open and closed arms was 45 and 5 luxes, respectively. Mice were placed in the central square facing one of the open arms and tested for 5 min. The percentage of time spent in the open arms was determined as a measure of anxiety. Animals that exit the maze during exploration were excluded. The 0-maze task was used to measure the acute effects of THC 30 min after the first administration. The elevated plus maze was performed to measure the effects of repeated THC 24 h after the last administration. For both tasks CD-1 mice were used (Suppl. Fig. 1b).

#### Barnes maze

Spatial learning and memory was assessed using the Barnes maze, as previously described (Gomis-González et al., 2021). The maze consists of a circular platform (90 cm in diameter) with 20 equally spaced holes through which mice may escape from a bright light (300 lx). Only one hole allows the escape to a dark/target box. Visual cues were placed surrounding the maze for navigational reference. Smart v3.0 software was used to control the video-tracking system. Briefly, mice were first habituated to the maze. In this phase, animals were placed in the centre of the maze for 10 sec covered by an opaque cylinder. After removal of the opaque cylinder, mice were gently guided to the target hole by surrounding them within a transparent cylinder so mice could see where the scape hole was located. Then, they were left inside the target box for 2 min and returned to the home cage. 1 h later, the first training phase was carried out. During training, each mouse performed 2 trials per day on 4 consecutive days. Each training trial started with the mouse placed in the centre of the Barnes maze covered by an opaque cylinder for 10 sec. Then, animals could explore the maze for 3 min. During this period, the time the mice spent to first visit the target hole was counted (primary latency). Each training trial ended when the mouse entered to the target box or after 3 min of exploration. When mice reached the target box, they stayed there for 1 min before returning to the home cage. When mice did not reach the target box within 3 min, the experimenter guided the mouse gently to the escape box within the transparent cylinder. Primary latency (time to reach the target box) was considered a measure of learning performance. For this task, mice were treated with vehicle or THC for 7 d and the habituation and training started 24h after the last administration. CD-1 animals were used (Suppl. Fig. 1b).

### Tissue preparation for immunofluorescence

BL-6 mice were perfused 24 h after the last administration of the sub-chronic treatment (7 d) with vehicle or THC 1 mg/kg (Suppl. Fig. 1c). Mice were deeply anesthetized by intraperitoneal injection (0.2 mL/10 g of body weight) of a mixture of ketamine (100 mg/kg) and xylazine (20 mg/kg) prior to intracardiac perfusion with 4 % paraformaldehyde in 0.1 M Na2HPO4/0.1 M NaH2PO4 buffer (phosphate buffer, PB), pH 7.5, delivered with a peristaltic pump at 19 mL/min flow for 3 min. Subsequently, brains were extracted and post-fixed in the same fixative solution for 24 h and transferred to a solution of 30 % sucrose in PB overnight at 4 °C. Coronal frozen sections (30 μm) of the dorsal hippocampus (coordinates relative to Bregma: – 1.22 mm to – 1.82 mm), basolateral amygdala (from Bregma: − 1.22 mm to − 1.82 mm) and the prelimbic prefrontal cortex (from Bregma: 1.98 mm to 1.54 mm) were obtained on a freezing microtome and stored in a solution of 5 % sucrose at 4 °C until used.

### Immunofluorescence

Free-floating brain slices were rinsed in PB, blocked in a solution containing 3 % donkey serum (DS) (Sigma-Aldrich) and 0.3 % Triton X-100 (T) in PB (DS-T-PB) at room temperature for 2 h, and incubated overnight in the same solution with the primary antibody to c-Fos (sc-7202, 1:1,000, rabbit, Santa Cruz Biotechnology) at 4 °C. The next day, after 3 rinses in PB, sections were incubated at room temperature with the secondary antibody AlexaFluor-555 goat anti-rabbit (1:1,000, Life Technologies, Thermo Fisher Scientific) in DS-T-PB for 2 h. After incubation, sections were rinsed and mounted immediately after onto glass slides coated with gelatin in Fluoromont-G with 4′,6-diamidino-2-phenylindole (DAPI) (Invitrogen, Thermo Fisher Scientific) as counterstaining.

### Image analysis

Immunostained brain sections were analysed with a ×10 objective using a Leica DMR microscope (Leica Microsystems) equipped with a digital camera Leica DFC 300FX (Leica Microsystems). Seven different brain subregions were analysed: prelimbic cortex (PL), basal amygdala (BA), central amygdala (CeA), lateral amygdala (LA), dentate gyrus (DG), Cornu Ammonis 1 (CA1) and 3 (CA3). For PL analysis, a 430-μm-sided square region of interest (ROI) was delimited for quantification. For the rest of regions, the DAPI signal was used as counterstaining to identify and delimitate each area for quantification. The images were processed using the ImageJ analysis software. c-Fos-positive neurons in each brain area were quantified using the automatic “particle counting” option with a fixed configuration that solely detected c-Fos-positive cell bodies matching common criteria of size and circularity. A fixed threshold interval was set to distinguish the c-Fos-positive nuclei from the background. In addition, all quantifications were individually checked for homogeneity by an expert observer blind to the experimental conditions. 4-6 representative brain sections of each mouse were quantified, and the average number of c-Fos-positive neurons was calculated for each mouse. The data are expressed as density: the mean number of c-Fos-positive cells per squared mm (n = 6 mice per experimental group). For the c-Fos data, the displayed images were flipped for orientation consistency and transformed to inverted grey scale. Network analyses were conducted computing correlations between activity in each structure within each group. Pearson r correlation values between brain-regions were calculated for each experimental group. Graph network analysis was performed in R (version 4.0.4) using the DescTools (version 0.99) package.

### Tissue preparation for in situ proximity ligation assay

CD-1 mice were perfused immediately after the test phase of NOR, 24 h after the last administration of the sub-chronic pre-treatment (7 d) with vehicle/ MDL 0.01mg/kg and treatment with vehicle/THC 1 mg/kg (Suppl. Fig. 1d). Mice were processed for intracardiac perfusion and brain removal as described above. Prior to sectioning, each brain was washed twice for 5 min with NaCl 0.9 % in PB (PBS). After briefly drying, brains were frozen in dry-ice cold methylbutane for 10 sec. Once frozen, coronal frozen sections (15 μm) of the dorsal hippocampus (coordinates relative to Bregma: – 1.22 mm to – 1.82 mm), and the prelimbic prefrontal cortex (from Bregma: 1.98 mm to 1.54 mm) were cut on a cryostat, mounted on super frost slides (Thermo Fisher) and stored at -80 °C until used.

### In situ proximity ligation assay

Heterodimers were detected using the Duolink II in situ proximity ligation assay (PLA) detection Kit (Sigma) and following the instructions of the supplier. To detect CB1R/5-HT2AR heterodimers, a mixture of equal amounts (1:100) of guinea pig anti-CB1R antibody (Frontier Science) and rabbit anti-5-HT2AR antibody (Neuromics) was used together with PLA probes detecting guinea pig or rabbit antibodies. Then, brain slices were processed for ligation and amplification with a Detection Reagent Red and were mounted using a mounting medium with DAPI. The samples were observed in a Leica SP2 confocal microscope (Leica Microsystems) equipped with an apochromatic 63X oil-immersion objective (1.4 numerical aperture), and a 405 nm and a 561 nm laser line. For each field of view, a stack of two channels (one per staining) and 6 to 10 Z stacks with a step size of 1 μm were acquired. Images were opened and processed with ImageJ confocal software (National Institutes of Health). Nuclei and red spots were counted on the maximum projections of each image stack. After getting the projection, each channel was processed individually. The nuclei were segmented by filtering with a median filter, subtracting the background, enhancing the contrast with the Contrast Limited Adaptive Histogram Equalization (CLAHE) plug-in, and finally applying a threshold to obtain the binary image and the regions of interest (ROIs) around each nucleus. Red spot images were also filtered and thresholded to obtain the binary images. Red spots were counted in each of the ROIs obtained in the nuclei images. The percentage of cells containing one or more red spot (positive cells) versus the total number of cells (blue nucleus) were determined using the Fiji package (http://pacific. mpi-cbg.de/), considering a total of 800–1,000 cells from 3 to 7 different fields within each brain region from the different mice per group.

### Hippocampal slice preparation for structural plasticity

Thy1-EGFP mice were perfused 24 h after the last administration of the sub-chronic treatment (7 d) with vehicle or THC 1 mg/kg (Suppl. Fig. 1c). Mice were processed for intracardiac perfusion and brain removal as described above 60 μm brain slices were obtained using a freezing microtome (LEICA SM 2000). Finally, slices were washed in 0.1 M PB, and mounted in glass slides with mowiol for subsequent confocal analysis.

### Confocal image acquisition and dendritic spine analysis

Images of dendritic spines from CA1 pyramidal neurons were obtain using a Leica TCS SP5 confocal microscope. Image stacks were of 0.13 μm step thickness. Dendritic spines counting and classification (mushroom, thin, stubby) was assessed through Neuronstudio program (Version 0.9.92; http://research.mssm.edu/cnic/tools-ns.html, CNIC, Mount Sinai School of Medicine) after deconvolution of the images by Huygens Essential software (Huygens compute engine 4.3.1p3 64b, https://svi.nl/HuygensSoftware). We analysed secondary and tertiary dendrites between 20 to 150 µm distance from soma. Spine analysis was carried out along 30 to 80 µm dendrite segment length. The parameters used for the counting and classification were as follow. Voxel dimensions X: 0.08 µm, Y: 0.08 µm, Z: 0.125 µm. For neurite tracing attach ratio: 1.3, Min Length: 3.0 and Discretion Ratio: 1.0. Dynamic and scattered sampling options were selected. For spine detection and classification, the program was trained with 300 examples of spines and analysis was then automatically performed. Clear erroneous detections were manually corrected by an expert observer blind to the experimental conditions.

### Hippocampal slice preparation for electrophysiology

Brain slices were prepared from BL-6 mice 24 h after the last of 7 consecutive daily doses of THC (1 mg/kg) or the vehicle (Suppl. Fig. 1c). Isoflurane was used to anesthetize mice, and euthanasia was performed by cervical dislocation followed by decapitation. The brain was removed and quickly introduced into modified cold, oxygenated (95% O_2_-5% CO_2_) artificial cerebrospinal fluid (m-aCSF) containing NaCl 109 mM, KCl 4.5 mM, MgCl_2_ 1 mM, CaCl_2_ 2.5 mM, NaH_2_PO_4_ 1.2 mM, NaHCO_3_ 35 mM, Glucose 11 mM, HEPES 20 mM, sodium ascorbate 0.4 mM. Transverse hippocampal slices (280 µm) were cut using a vibrating tissue slicer (Leica VT1200S, Leica Biosystems) while submerged in oxygenated m-aCSF. They were then transferred to continuously oxygenated aCSF, maintained at 35° C for 20 min. Following this, they were maintained at room temperature in oxygenated aCSF for at least 40 min prior to recording. A maximum of five slices per animal per day were used for electrophysiological recordings.

### Electrophysiological recordings

Extracellular field excitatory postsynaptic potentials (fEPSPs) were recorded in area CA1 of the hippocampal slices using previously described techniques (Lecca et al., 2019). The aCSF used for the recordings contained: NaCl 126 mM, KCl 3 mM, MgCl2 1 mM, CaCl2 2.4 mM, NaH2PO4 1.2 mM, NaHCO3 26 mM, glucose 11 mM. To limit the influence of endogenous adenosine on fEPSPs, caffeine (50 µM) was included in the aCSF. Hippocampal slices were submerged in a recording chamber (RC-26, Warner Instruments, Hamden, CT, USA), and continuously perfused with aCSF (2 mL/min) using a peristaltic pump (Cole-Parmer Instruments, Vernon Hills, IL, USA). Bath temperature was maintained at 30–32 °C by passing the aCSF through an in-line heater (TC324-C and SH27-B, Warner Instruments). Borosilicate glass electrodes (1.5mm o.d. × 0.86 mm i.d., Sutter Instruments, Novato, CA, USA) were fabricated using a horizontal puller (P-97, Sutter Instruments) and filled with aCSF. Electrodes were connected to the headstage of an AC amplifier (Model 1800, A-M Systems, Sequim, WA, USA). The tip of a bipolar-stimulating electrode consisting of twisted formvar-insulated nichrome wire (50 μm diameter; A-M Systems), connected to a constant current stimulus isolation unit (DS3, Digitimer LLC, Ft. Lauderdale, FL), was positioned in hippocampal area CA3 to activate Schaffer collateral axons forming synapses on CA1 pyramidal neuron dendrites. The recording electrode was positioned in CA1 stratum radiatum using a manual micromanipulator, and gradually lowered while monitoring fEPSPs, so that the largest fEPSP for a given stimulus intensity was obtained. Recording electrodes were then fixed at this depth within the slice and input–output (I/O) curves relating fEPSP amplitude to stimulus intensity (20–200 μA, 0.1 ms) were generated. Baseline fEPSP responses of 30–50% of maximum were then obtained using a stimulus intensity determined from the I/O relationship (which was not statistically different among groups, data not shown). Following establishment of a stable baseline period of 15 min, theta burst stimulation (TBS) was applied in order to induce long-term potentiation (TBS-LTP). The TBS protocol consisted of 10 bursts of 5 stimuli delivered at 100 Hz, separated by 200 msec. Episodes were repeated 4 times at 5 s intervals. The stimulation, data acquisition and signal analyses were performed online using an A/D board (PCIe-6321, National Instruments, Austin, TX, USA) and WinLTP software (https://www.winltp.com) (Anderson and Collingridge, 2007).

### RNA extraction and reverse transcription

Brain samples were obtained from CD-1 mice 24 h after the last administration of the sub-chronic treatment (7 d) with vehicle or THC 1 mg/kg (Suppl. Fig. 1c). Hippocampal tissues were rapidly dissected and stored at −80 °C un8l used. Isola8on of total RNA was performed using the RNeasy Mini kit (QIAGEN) according to the manufacturer’s instructions. Total RNA concentration was measured using a NanoDrop spectrophotometer (Thermo Fisher Scientific). Reverse transcription was performed with 10 μL of total RNA from each animal to produce cDNA in a 20-μL reaction using the High Capacity cDNA Reverse Transcription kit (Applied Biosystems) according to the manufacturer’s instructions. The cDNAs from brain tissues was diluted to a final concentration of 10 ng/μL and stored at −20 °C un8l use.

### Quantitative real-time PCR analysis

Real-time PCR was carried out in a 10 μL reaction using SYBR Green PCR Master Mix (Roche) using 1ng/μL of cDNA according to the manufacturer’s protocol with a QuantStudio 12 K Flex Real-Time PCR System (Applied Biosystems). The following primers specific for mouse were used:

Bdnf (BDNF) 5’-CAGGTGAGAAGAGTGATGACC-3’ 5’-ATTCACGCTCTCCAGAGTCCC-3’

Ngf (NGF) 5’-CAAGGACGCAGTTTCTATACTG-3’ 5’-CTTCAGGGACAGAGTCTCCTTCT-3’

Actb (β-actin) 5’-CGTGAAAAGATGACCCAGATCA-3’ 5’-CACAGCCTGGATGGCTACGT-3’

All primers were used at 10 μM. Quantification was performed by using the comparative CT Method (ΔΔCT Method). All the samples were tested in triplicate and the relative expression values were normalized to the expression value of β-actin. The fold change was calculated using the eq. 2^(–ΔΔCt)^

### Statistical analysis

Comparisons between groups were performed by Student t test or one-way or two-way analysis of variance (ANOVA) for multiple-group comparisons, depending on the factors involved. Post hoc comparisons were performed by Bonferroni test only when significant main effect of one-way ANOVA or significant effect of factors or interaction between factors of two-way ANOVA were revealed or significant repeated-measure ANOVA. For morphology of spine density, Post hoc comparisons were performed by Newman-Keuls test. The statistical analysis and the artwork was performed using GraphPad Prism 7. All results were expressed as mean ± s.e.m. Differences were considered statistically significant when p <0.05.

## Results

### Repeated exposure to a THC dose without acute amnesic-like effects produces deficits in hippocampal long-term memory

A single dose of THC at 10 mg/kg or 3 mg/kg produces significant novel object-recognition (NOR) memory impairment (Busquets-Garcia et al., 2018), with no signs of tolerance after 7 d of repeated administration (Puighermanal et al., 2013). As many cannabis preparations used by humans can contain lower doses of THC, we wondered whether a dose of THC that does not acutely affect memory performance could eventually affect memory after repeated exposure (Fig. 1). We used a dose of THC (1 mg/kg; THC 1) that, as previously described, did not affect NOR memory when administered after the familiarization phase (Puighermanal et al., 2009). We administered THC 1 or vehicle to mice after the familiarization phase in a consecutive version of the NOR memory test. In this approach, the memory test phase of one day is equivalent to the familiarization phase for the next day, as previously described (Puighermanal et al., 2013) . Such repeated evaluation of NOR memory allowed detection of the deleterious effect of repeated THC 1 on memory consolidation after 5 administrations (Fig. 1a). After withdrawal of THC 1, NOR memory performance gradually improved, reaching control values after 2 d (Fig. 1a).

**Figure 1.**
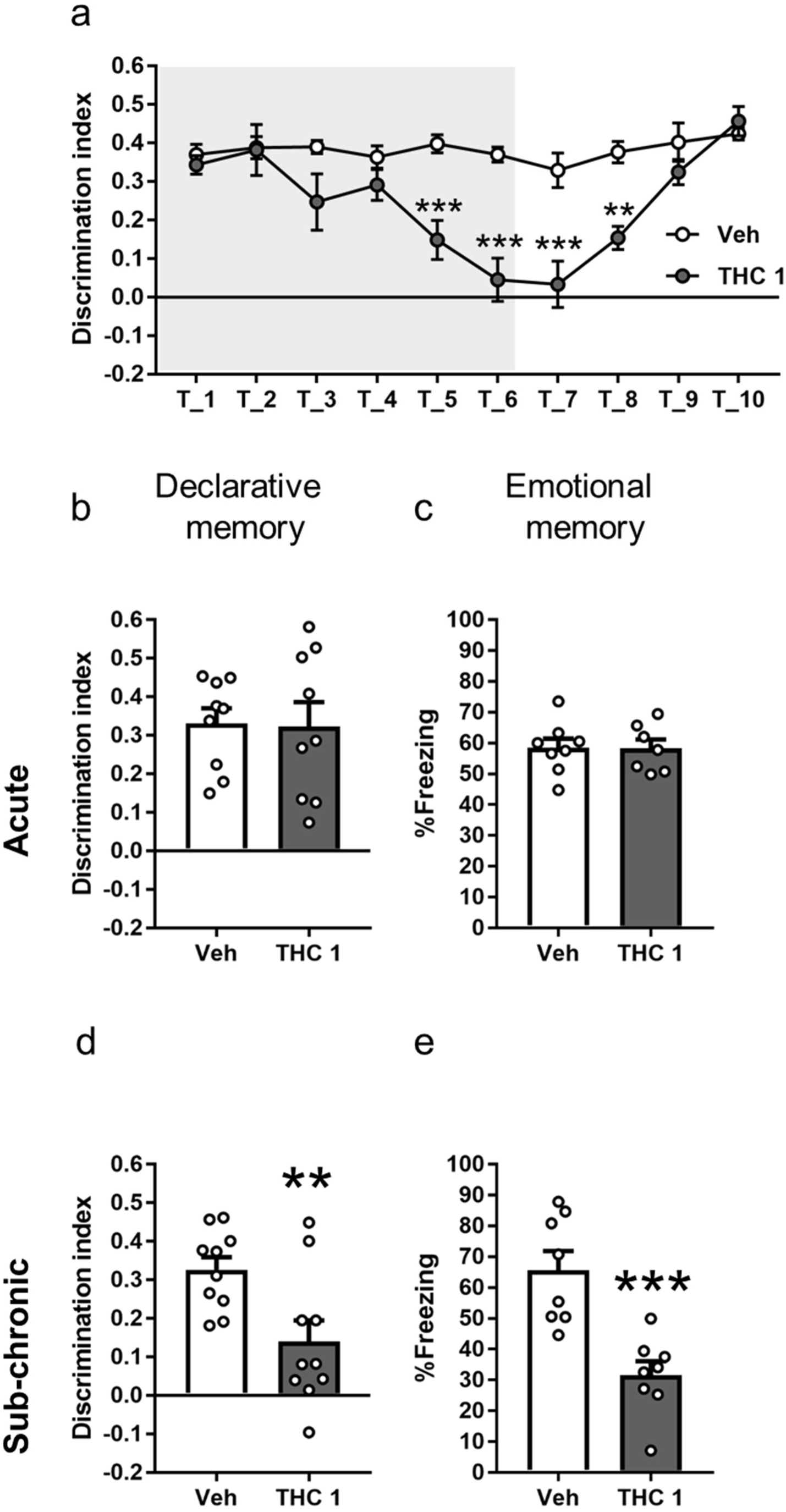
Repeated exposure to a sub-effective low dose of THC (1 mg/kg, THC 1) produces impairments in non-emotional and emotional memory in mice. (a) Consecutive NOR memory test in mice receiving THC 1 mg/kg (THC 1) or vehicle (Veh) on 6 consecutive days (grey area). Memory impairment was detected after 5 administrations, and animals treated with THC totally recovered memory performance two/three days after THC 1 withdrawal (Veh n = 5-10, THC 1 n = 8-10). Statistical significance was calculated by Bonferroni post hoc test following two-way ANOVA. ** p< 0.01, *** p< 0.001 (Veh vs THC 1). A single dose of low-dose THC (1mg/kg) did not produce memory impairment in (b) NOR memory (Veh n = 9, THC 1 n = 9) or in (c) Trace FC (Veh n = 8, THC 1 n = 7). However, after a repeated exposure THC 1 produced significant memory impairments detected in (d) NOR task (Veh n = 10, THC 1 n = 10) and in (e) Trace FC (Veh n = 8, THC 1 n = 8) when these tasks were performed 24 h after the last THC 1 administration. Statistical significance was calculated by Student’s t-test. ** p< 0.01, *** p< 0.001. Data are expressed as mean ± S.E.M.

Considering this result, we decided to study the effects of an acute vs. a sub-chronic (7 d) administration of THC (1 mg/kg) on both non-emotional and emotional hippocampal memory and spatial working memory. In agreement with the observation in Fig. 1a, a single administration of THC 1 after familiarization/training affected neither NOR memory (Fig. 1b) nor trace fear conditioning (Trace FC) performance (Fig. 1c). Furthermore, no effect on the exploratory behaviour was observed in the NOR task (Suppl. Fig. 2a).

Next, in the cohort previously used for studying NOR memory, mice were injected with THC 1 every day for 6 additional days without behavioural assessment, and the seventh administration of THC 1 was delivered 20 min after the familiarization phase for the NOR memory test (Suppl. Fig. 1a). Then, NOR memory was assessed the next day. Under these experimental conditions, mice exposed to sub-chronic THC 1 showed, as expected, a significant decrease in discrimination index (Fig. 1d) with no effect on the exploratory behaviour (Suppl. Fig. 2b).

Using another cohort of mice, we assessed the effect of 7 doses of THC 1 in the Trace FC paradigm. Emotional hippocampal memory was severely impaired after repeated THC 1 administration, similar to what we observed for non-emotional hippocampal memory in the NOR task (Fig. 1e). However, when spatial working memory was tested in another cohort of mice using the Y-maze, no alterations in performance or exploratory behaviour were observed with a single administration, nor following 7 d of sub-chronic exposure to THC 1 (Suppl. Fig. 3).

We additionally assessed anxiety-like behaviour in the same conditions of acute and sub-chronic administration of THC 1, since the effects of THC are biphasic in this behavioural trait (Puighermanal et al., 2013), but we did not observe any significant effects following either acute or sub-chronic exposure to THC 1 on either the 0-maze (Suppl. Fig. 4a) and the elevated plus maze (Suppl. Fig. 4b), respectively.

### Cognitive outcome of repeated THC 1 exposure varies according to the behavioural test

We further assessed the outcome of repeated THC 1 exposure beyond drug withdrawal. In order to study spatial learning, we used the Barnes maze (Paul et al., 2009) . Mice previously exposed to THC 1 or vehicle were trained for 4 consecutive days after treatment withdrawal. We found that spatial learning was preserved under our experimental conditions, since both groups showed similar primary latency values to find the escape hole in every test (Fig. 2a). We then analysed the outcome of THC 1 exposure using the NOR and FC tests and performing the training sessions on the next day after treatment withdrawal. To investigate NOR deficits, mice were treated for 7 d and 24 h after the last administration they were subjected to the NOR familiarization phase. The following day, 48 h after the last administration of THC 1, memory was evaluated. Using this protocol, a significant deficit in NOR memory was observed in mice previously exposed to THC 1 (Fig. 2b).

**Figure 2.**
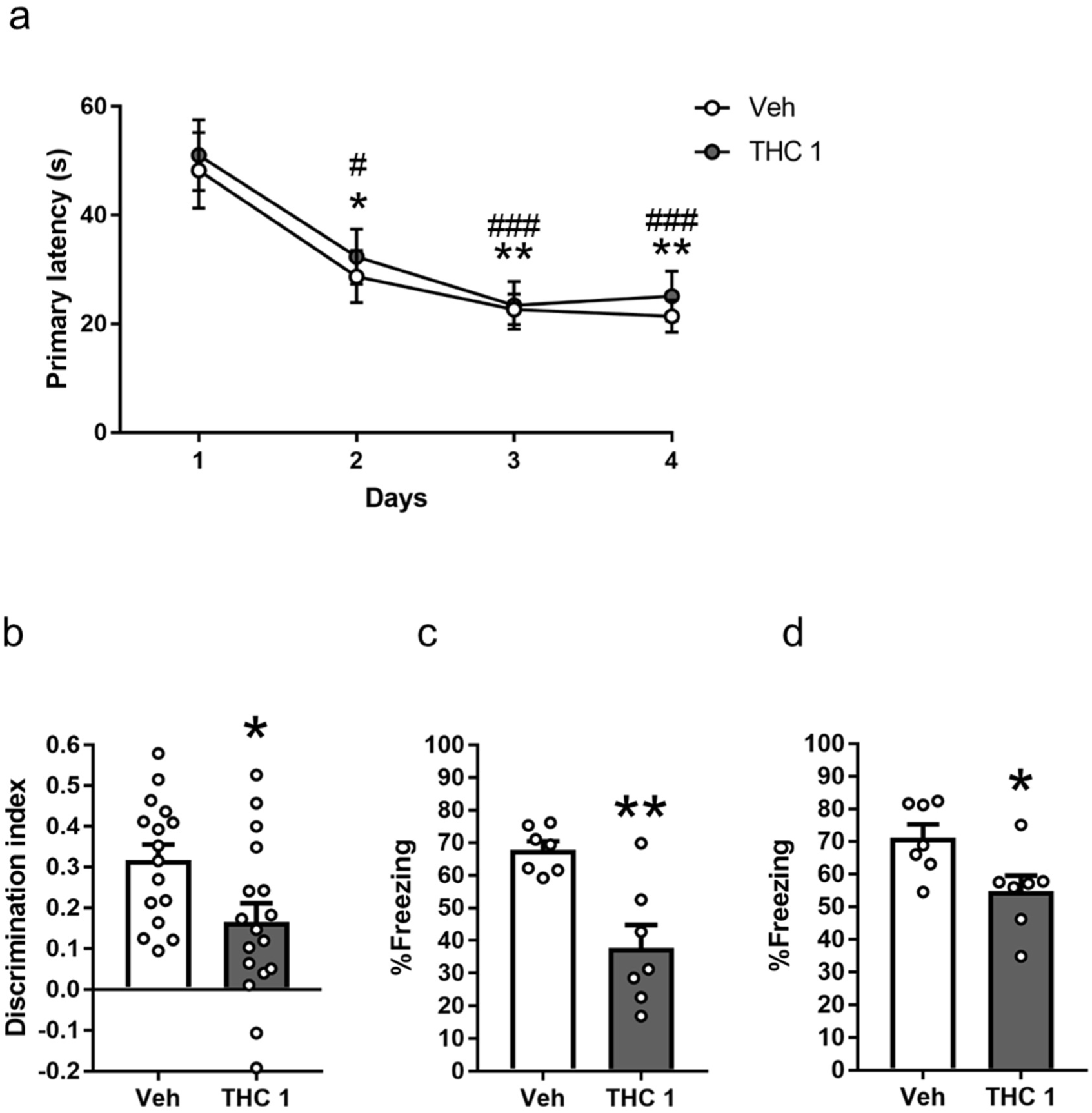
Cognitive outcome of THC 1 exposure varies according to the behavioural test. (a) Spatial learning, as assessed in the Barnes maze task, did not show differences between treatment conditions as demonstrated by the similar decrease in primary latency over the training days (Veh n = 18, THC 1 n = 16). Statistical significance was calculated by Bonferroni post hoc test following two-way ANOVA repeated-measures. * p < 0.05, ** p < 0.01 (vs Day 1 Veh). # p < 0.05, ### p < 0.001 (vs Day 1 THC 1). (b) NOR memory task performed 48 h after the last THC 1 administration showed significant alterations in non-emotional memory (Veh n = 16, THC 1 n = 17). For the emotional memory, sub-chronic exposure with THC 1 mg/kg produced a memory impairment in (c) Trace FC performed 48h after the last administration (Veh n=7, THC 1 n=7) and in (d) Context FC performed 72h after the last administration in the same cohort of mice (Veh n=7, THC 1 n=7). Statistical significance was calculated by Student’s t-test. * p< 0.05, ** p< 0.01. Data are expressed as mean ± S.E.M.

Notably, when Trace FC training was performed the next day after drug withdrawal, the THC 1 exposed group demonstrated a significant memory impairment when mice were exposed to the conditioned stimulus (Fig. 2c), but also when treated mice were exposed to the conditioning box the next day (72 h after the last THC administration) (Fig. 2d) Together, specific alterations in learning and memory performance were derived from THC 1 exposure that extended beyond the treatment period.

### Repeated THC 1 exposure produces alterations in functional and structural synaptic plasticity

Alterations in functional and structural synaptic plasticity in the hippocampus have been described after chronic exposure to high doses of THC (10 mg/kg) (Hoffman et al., 2007; Chen et al., 2013). In order to determine whether synaptic plasticity is also altered by chronic, low-dose THC, we examined LTP in slices obtained from mice following 7 d THC 1 treatment. Theta burst stimulation produced LTP of the fEPSP in hippocampal slices from both vehicle and THC 1-treated mice. However, the magnitude of LTP observed after theta burst stimulation was significantly decreased in the THC 1 group in the last 10 min of the recording session (Fig. 3a; Fig. 3b).

**Figure 3.**
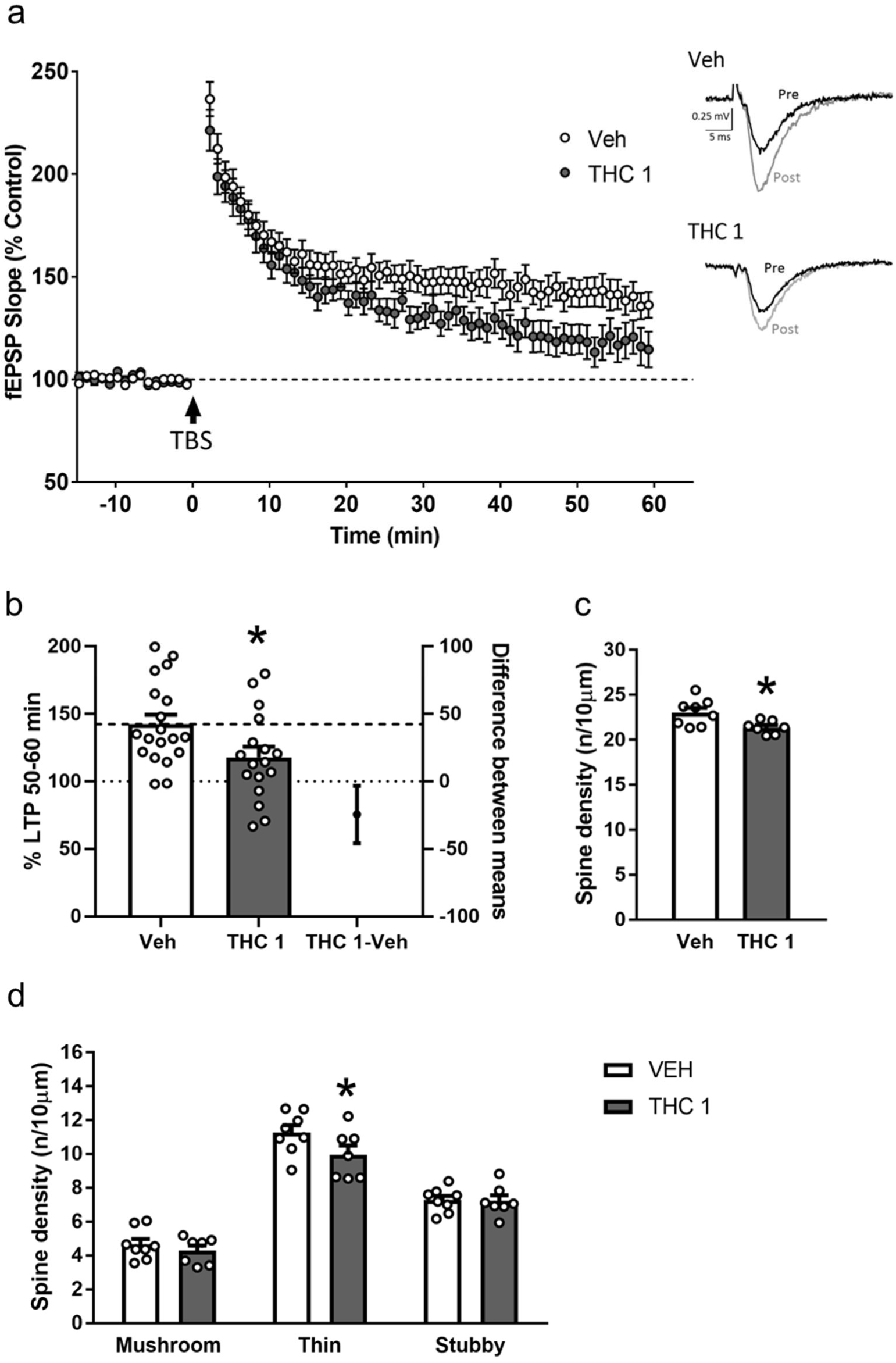
Sub-chronic low-dose THC inhibits synaptic plasticity in CA1 region of the hippocampus and decreased total spine density. (a) The average time courses of the change in the slope of the fEPSP in hippocampal slices from mice treated for 7 d with Veh or THC 1. After stimulation, slices from THC 1-treated mice showed a small but significant decrease in LTP at Schaffer collateral synapses (Veh n = 19 slices/4 mice THC 1 n = 17 slices/4 mice). (b) Average fEPSP slope values obtained between 50 min and 60 min after theta burst stimulation (TBS) showing a significant decrease in THC 1 treated slices (Veh n = 19 slices/4 mice THC 1 n= 17 slices/4 mice). Statistical significance was calculated by Student’s t-test. * p< 0.05. (c) Sub-chronic treatment with THC 1 decreased CA1 pyramidal neuron overall spine density (Veh n = 8, THC 1 n = 7). Statistical significance was calculated by Student’s t-test. * p< 0.05. (d) Spine morphology analysis showed a significant decrease in thin spine density to (Veh n = 8, THC 1 n = 7). Statistical significance was calculated by Newman-Keuls post hoc test following two-way ANOVA. * p< 0.05 (Veh vs THC 1). Data are expressed as mean ± S.E.M.

Thy1-EGFP transgenic mice were used to study spine density and morphology in the CA1 pyramidal neurons expressing the fluorescent protein EGFP. As above, animals were treated with THC 1 or vehicle for 7 d and 24 h after the last injection histological samples were obtained and CA1 pyramidal dendritic spines were analysed. A significant decrease in total dendritic spine density was observed after THC 1 exposure compared to vehicle-treated mice (Fig. 3c). When the spine structure was analysed, we found a significant decrease in thin spines in the THC 1 treated mice (Fig. 3d).

These alterations in functional and structural plasticity were not accompanied by changes in RNA levels for two of the most studied neurotrophins BDNF and NGF (Suppl. Fig. 5).

### Sub-chronic administration of THC 1 altered overall c-Fos expression and correlation patterns between memory-related brain regions

Acute THC exposure also altered functional connectivity as measured by c-Fos expression analysis between brain regions involved in reward and stress responses (Ruiz et al., 2021). We hypothesized that, after the sub-chronic exposure to THC 1, neuronal activity, measured by c-Fos expression analysis, and connectivity, measured by the correlation analysis of c-Fos expression between memory-related brain regions, could be disturbed. We performed the analysis of c-Fos expression the next day after treatment withdrawal to circumvent any acute effect of THC. Therefore, we treated mice for 7 d with THC 1 and 24 h after the last administration brain samples were obtained and c-Fos expression was assayed in seven memory-related brain regions including amygdala (BA, CeA and LA), hippocampal (DG, CA3 and CA1) and prefrontal (PL) regions. An overall enhanced c-Fos expression was detected in the regions analysed (Fig. 4a), with a significant increase in hippocampal CA1 region (Fig. 4b).

**Figure 4.**
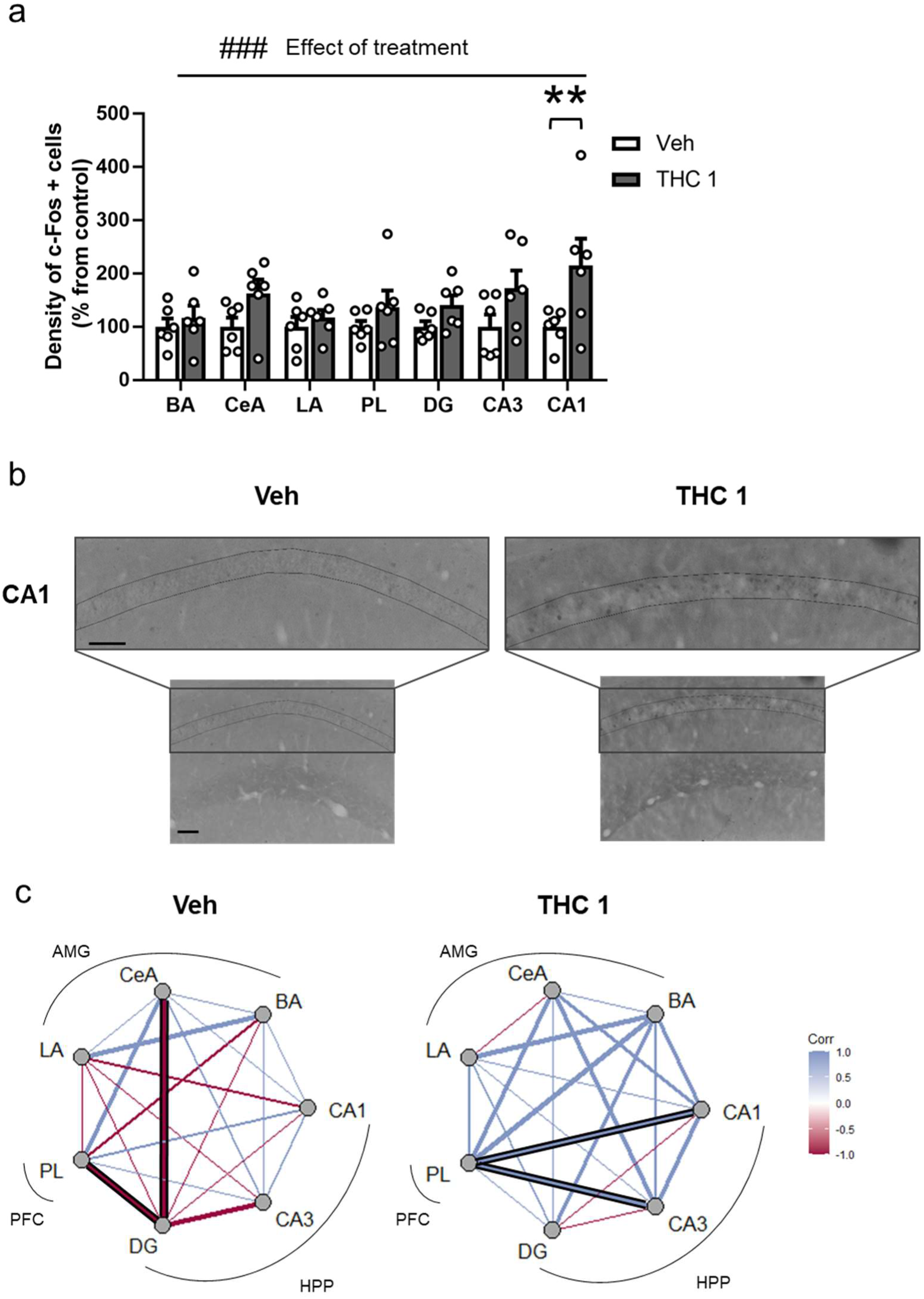
Sub-chronic THC 1 mg/kg exposure enhances overall c-Fos expression and modifies c-Fos correlation pattern between brain regions. (a) Quantification of c-Fos immunudetection density in selected brain regions after THC 1 of vehicle (Veh) exposure (Veh n = 6, THC 1 n = 6). Statistical significance was calculated by Bonferroni post hoc test following two-way ANOVA. ### p < 0.001 (main treatment effect), ** p < 0.01 (post-hoc). Data are expressed as mean ± S.E.M. (b) Representative images of c-Fos expression in Veh-and THC 1-treated animals. Scale bar= 75 µm. (c) Circos plot analysis of c-Fos expression in different brain regions after sub-chronic THC administration. Network graphs of c-Fos expression based on the Pearson coefficient in Veh and THC 1 between different brain areas, including dentate gyrus (DG), cornu ammonis 1 (CA1), cornu ammonis 3 (CA3), prelimbic cortex (PL), basal amygdala (BA), lateral amygdala (LA) and central amygdala (CeA). Colours depict correlation indexes. Significant correlations are delimited with black lines.

Subsequent connectivity analysis between control and treated mice revealed an alteration in network connectivity after sub-chronic THC 1 administration. In the vehicle-treated group, the c-Fos expression on most of the brain regions analysed were negatively correlated. Specially, a significant functional de-coupling was observed between DG-CeA and DG-PL (Fig. 4c). After THC 1 treatment, this connectivity map was strongly altered and the activity of most of the brain regions analysed were positively correlated, indicating an increase in connectivity following THC 1 treatment. Importantly, a significant functional coupling was observed between the PL-CA1 and PL-CA3 regions (Fig. 4c).

### 5-HT2AR blockade prevents THC 1 deficits in non-emotional and emotional hippocampal long-term memory

Since the serotonergic system, and more specially 5-HT2ARs, have been implicated in non-emotional amnesic-like effects of acute THC (Viñals et al., 2015), we hypothesized that the memory impairment that we observed after low-dose THC could be also mediated by the serotonergic system. We pre-treated mice with MDL 100,097 (MDL), a 5-HT2A antagonist, before each THC 1 administration and analysed the effects of this on non-emotional and emotional memory in different cohorts of mice. In agreement with previous results, a single acute administration with THC 1 or with MDL did not alter object-recognition memory (Suppl. Fig. 6a). However, the amnestic effects of repeated THC 1 were completely prevented by pre-treatment with MDL. (Fig. 5a). To investigate whether a similar involvement of 5-HT2ARs was observed in the emotional memory deficit produced by THC 1, two different cohort of mice were used. In the first, we confirmed that acute THC 1 and MDL did not affect Trace FC memory in the same conditions used for NOR task (Suppl. Fig. 6b). The second cohort of mice received repeated THC 1 with MDL pre-treatment, and we found that the emotional memory impairment produced by sub-chronic treatment with THC 1 was totally by the 5-HT2AR antagonist (Fig. 5b).

**Figure 5.**
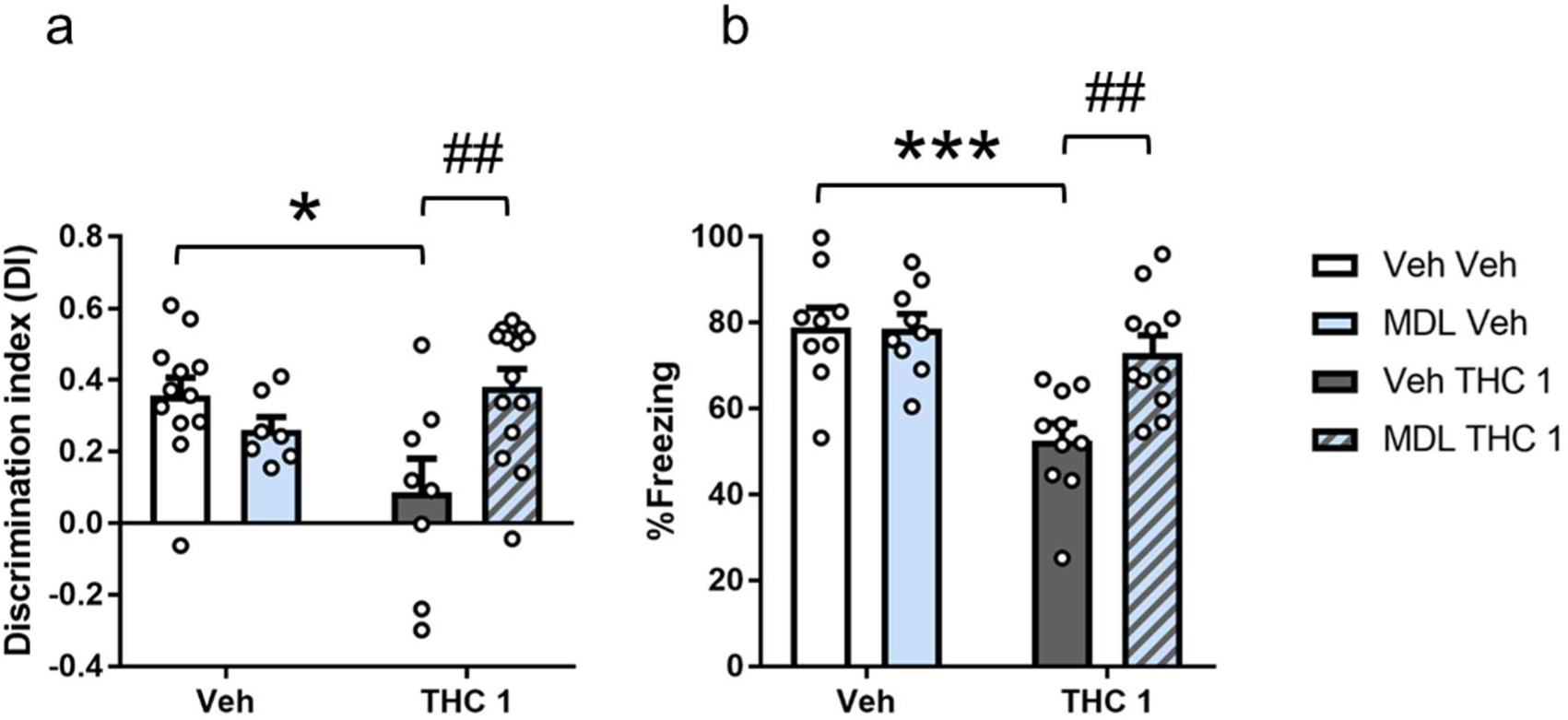
Memory impairment produced by sub-chronic THC 1 mg/kg administration is prevented with 5-HT2AR blockade. A sub-chronic pre-treatment with MDL 0.01 mg/kg prevented THC 1 amnesic-like effects in (a) declarative memory measured in NOR task (Veh Veh n = 12, MDL Veh n = 7, Veh THC 1 n = 8, MDL THC 1 n = 14) and (b) emotional memory measured in Trace FC (Veh Veh n = 9, MDL Veh n = 9, Veh THC 1 n = 10, MDL THC 1 n = 11). Statistical significance was calculated by Bonferroni post hoc test following two-way ANOVA. * p < 0.05, *** p < 0.001 (Veh Veh vs Veh THC 1). ## p < 0.01 (Veh THC 1 vs MDL THC 1). Data are expressed as mean ± S.E.M.

### Sub-chronic exposure to THC 1 enhances the expression CB1R/5-HT2AR heterodimers

The involvement of 5-HT2ARs in the memory impairment caused by repeated THC 1 suggested that CB1R/5-HT2AR heterodimers might be involved. Indeed, enhanced levels of CB1R/5-HT2AR heterodimers have been detected in cannabis consumers (Galindo et al., 2018; Guinart et al., 2020). Therefore, the expression of these heterodimers after repeated THC 1 exposure was measured. Thus, a cohort of mice received THC 1 treatment alone, and another cohort received the MDL pre-treatment, as detailed above. Then, brain tissue was collected after the performance in the NOR test. We observed that THC 1 treatment produced a significant enhancement in CB1R/5-HT2AR heterodimers in the hippocampus and that MDL pre-treatment prevent the increase in heterodimers (Fig. 6a). Notably, the enhanced appearance of the heterodimers was not observed in the prefrontal cortex, another brain region where both receptors are expressed (Suppl. Fig 7a). In addition, a correlation analysis was also performed between the NOR task discrimination index and heterodimer expression considering the entire population (n = 16) and we observed a significant negative correlation between the memory performance and the levels of CB1R/5- HT2AR heterodimers (Fig 6b), supporting the involvement of enhancement in those heterodimers in cognitive deficits.

**Figure 6.**
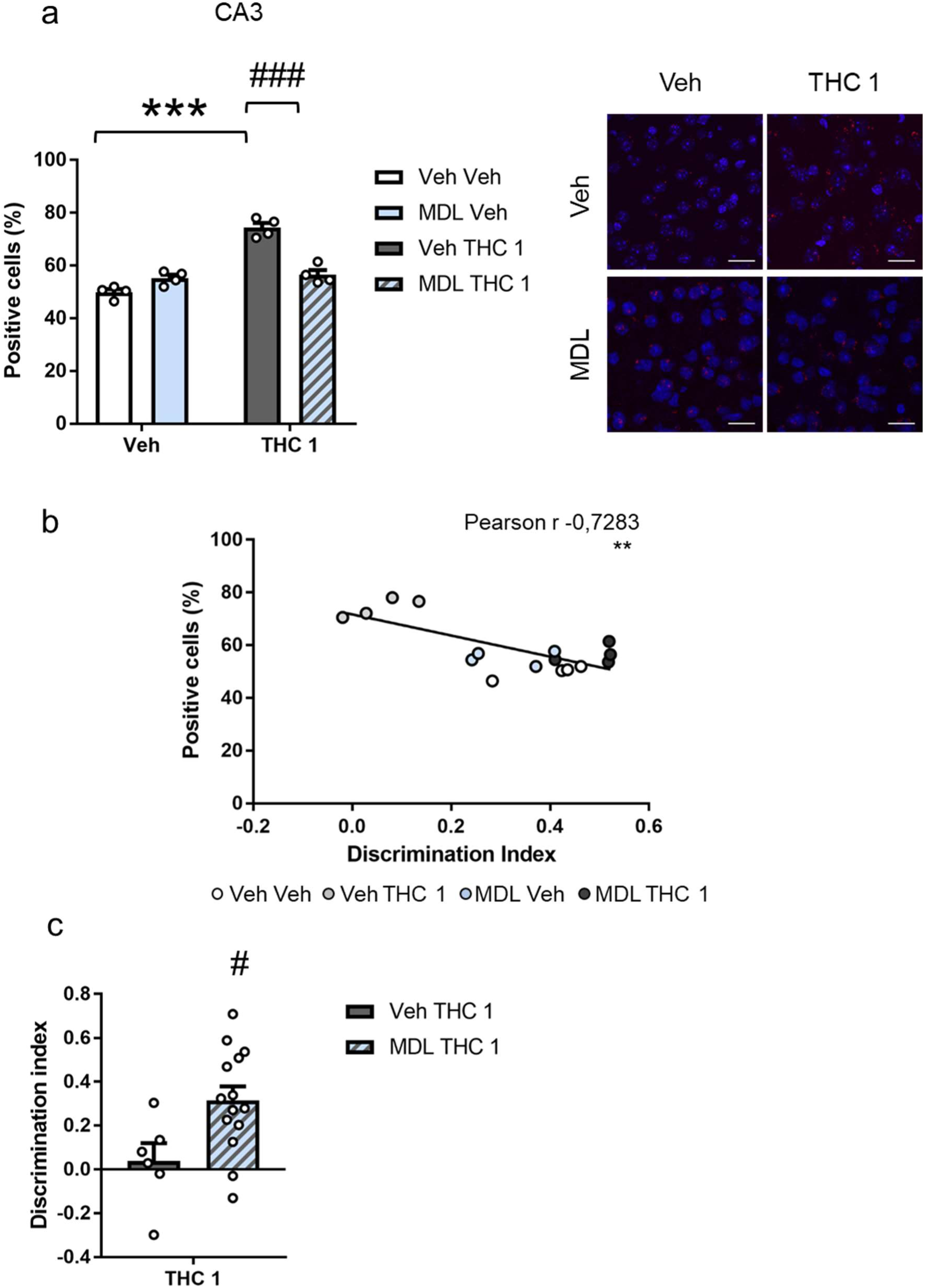
CB1R/5-HT2AR heterodimer detection is enhanced after repeated THC 1 mg/kg exposure. (a) THC 1 exposure significantly enhanced the detection of cells showing CB1R/5-HT2AR heterodimers, and sub-chronic pre-treatment with MDL prevent the up-regulation of heterodimers formed between CB1 and 5-HT2A receptor after sub-chronic THC 1 treatment in the CA3 region of the hippocampus (Veh Veh n=4, MDL Veh n=4, Veh THC 1 n=4, MDL THC 1 n=4) (see Suppl. Figure 1d for treatment schedule) Representative images of the four groups, scale bar=20 µm. Statistical significance was calculated by Bonferroni post hoc test following two-way ANOVA. *** p < 0.001 (Veh Veh vs Veh THC 1). ### p < 0.001 (Veh THC 1 vs MDL THC 1). (b) A significant negative correlation between the memory indez and the levels of CB1R/5-HT2AR heterodimers was observed (n=16). Statistical significance was calculated by Pearson r. ** p < 0.01. (c) Acute MDL administration before the seventh administration of THC 1 also prevented THC 1 memory deficits in NOR task (Veh THC 1 n = 6, MDL THC 1 n = 14). Statistical significance was calculated by Student’s t-test. # p < 0.05 (Veh THC 1 vs MDL THC 1). Data are expressed as mean ± S.E.M.

To assess whether such enhanced expression of the CB1R/5-HT2AR heterodimers in the hippocampus was responsible for the amnesic-like effects of THC 1 treatment, we pre-treated with MDL acutely. In this case, all mice in the experiment were treated with THC 1 and only half of them received MDL 20 min prior the last THC 1 administration, coinciding with the familiarization phase of NOR task. To avoid possible bias between mice in the two groups the acute effect of THC 1 was analysed in all mice of the cohort to discard differences in the NOR task (Suppl. Fig.6c). We observed that administration of MDL on the last day of THC 1 treatment can prevent the amnesic-like effects of THC (Fig. 6c), supporting the involvement of enhanced expression of CB1R/5-HT2AR heterodimers by repeated THC 1 on cognitive performance.

## Discussion

In this study we describe the deleterious effect of a repeated low-dose of THC on memory performance revealing the CB1R/5-HT2AR heterodimers as significant substrates for such memory deficit.

We first assessed the effects of acute low-dose THC (1 mg/kg) on performance in different memory dimensions, including declarative non-emotional, emotional, and working memory. In line with previous findings (Puighermanal et al., 2009), this acute low-dose of THC did not affect any of these measures of memory. By analysing the effect of THC 1 in the repeated NOR memory paradigm, we observed that THC 1 developed a transient sensitizing effect for memory disruption. This is a somewhat surprising result, since repeated THC exposure produces a rapid tolerance to most of its pharmacological effects such as hypothermia, analgesia, or even anxiety-like effects (Puighermanal et al., 2013) at higher doses, without developing tolerance to the amnesic-like effect. We next determined whether repeated administration of this low dose THC would affect different types of memory. A significant memory impairment was observed in the NOR task and in emotional memory after repeated THC. This is consistent with reports of disrupted hippocampal-dependent NOR with acute higher doses of THC (Puighermanal et al., 2009, 2013; Viñals et al., 2015), and could be interpreted as caused by the development of sensitization. Regarding the disturbances on emotional memory after THC exposure, we firstly described them in a specific modality of fear conditioning, Trace FC, but this result is reminiscent to the previous described effect in another hippocampal-dependent emotional task, the Context FC, of acute high dose THC (Puighermanal et al., 2009) and other cannabinoid agonist (Maćkowiak et al., 2009). However, the repeated exposure to THC 1 did not affect working memory in our experimental conditions. This result is in contrast with the amnesic-like effects observed also in murine models after acute and repeated exposure to high doses of THC in working-memory (Varvel et al., 2001; Han et al., 2012). These results could indicate an independent modulation by low-dose THC on declarative and working memory which could be explored in future studies.

As observed in the repeated NOR paradigm, the deleterious effect of THC 1 treatment was evident when the training phase of the memory tests were performed 24 h after the last THC administration, likely indicating brain alterations that would go beyond THC presence, as THC is described to be no longer detected in the brain 24 h after repeated treatment with a THC (10 mg/kg) (Hoffman et al, 2007). The memory disruption after THC 1 withdrawal is in concordance with previous finding for object-recognition memory (Puighermanal et al., 2013) where significant amnesic-like effects of repeated THC (10 mg/kg) administration last for 5 d after treatment withdrawal. Taking into account the previous described effects of higher doses of THC and cannabinoids on spatial learning assessed using the Morris water maze (Cha et al., 2006), we decided to evaluate if our experimental conditions with repeated low-dose THC could also affect this type of memory on a similar paradigm that also measure spatial learning, the Barnes maze (Gawel et al., 2019). We found that mice receiving repeated low-dose THC treatment showed similar spatial learning to the vehicle-treated group. This result is in accordance with a previous study in which they observed that very high doses of THC (100 mg/kg) are needed to produce a spatial learning impairment compared to other pharmacological effects of THC (Varvel et al., 2001).

At the cellular and molecular level, we focused our analysis on alterations previously described for higher amnesic doses of THC. In particular, we examined the effects of repeated low-dose THC on synaptic plasticity using both functional and structural measures in the hippocampal CA1 region. We found that 1 d after repeated low-dose THC there was a significant reduction in theta-burst LTP at Schaffer collateral CA1 pyramidal neuron synapses. This observation is in agreement with previous findings where repeated treatment with a higher dose of THC (10 mg/kg) was found to block LTP at these synapses (Hoffman et al., 2007). However, this same study reported that LTP was not impaired 1 d after a single treatment with this same concentration of THC (10 mg/kg) (Hoffman et al., 2007). An examination of potential structural changes in the hippocampus showed a significant decrease in total dendritic spine density in CA1 pyramidal neurons following low-dose repeated THC. This is somewhat similar to a previously reported decrease in CA1 dendritic spine density observed after repeated treatment with a higher dose of THC (10 mg/kg) (Chen et al., 2013), but opposite to changes observed following low dose THC in aged mice (Bilkei-Gorzo et al., 2017). In further analysis we stratified the structural results based on different dendritic spine morphologies, and we observed that the decrease was most significant in thin spine density, which would indicate a reduction in plasticity potential of brain circuits (Chidambaram et al., 2019). Since neurotrophins are signalling molecules that can modify both synaptic transmission and structure in the hippocampus (Gibon and Barker, 2017) we decided to study the expression of two of the major neurotrophins in brain, BDNF and NGF at the mRNA level. No changes in these trophic factors were observed in the hippocampus, indicating that the impairment in synaptic plasticity produced by THC is not likely to be associated with modifications in the expression of these genes.

We also explored whether brain tissues would be affected with regard to their c-Fos responsiveness. Acute high and low doses of THC (10, 5 and 0.5 mg/kg) were previously found to induce the expression of the immediate-early gene c-Fos in diverse brain regions (Valjent et al., 2002; Flores et al., 2016; Ruiz et al., 2021). However, the effects of repeated low-dose THC on c-Fos expression has not yet been determined. We focused our analysis of c-Fos expression in the next day after THC withdrawal, since there is a clear memory deficit and clear alterations in synaptic plasticity, and to avoid the acute effect of the THC injection. A significant effect of THC exposure of c-Fos was observed in memory-related brain regions, including the CA1 subregion of the hippocampus, where we observed a decreased in spine density after THC exposure. Since c-Fos expression can also be used to assess the connectivity between regions, and previous findings pointed to altered functional connectivity in reward and stress networks after acute THC exposure (Ruiz et al., 2021), we examined the connectivity of memory-related brain regions. In vehicle-treated mice c-Fos expression among brain regions was negatively correlated. However, following the THC 1 treatment connectivity among brain regions was positively correlated. Specifically, c-Fos activity in prelimbic, CA1 and CA3 hippocampus was correlated, and this is in clear agreement with the previous study showing increased overall global connectivity and increased co-activation of PL and the hippocampal CA1–3 subregion in rats following single high-dose THC exposure (Ruiz et al., 2021). One of the limitations of our study is that all behavioural, molecular, pharmacological, and biochemical determinations have been performed only with male mice. However, sex-dependent differences in THC effects on all these levels have been described (Ruiz et al., 2021).

The serotonergic system has been implicated in several THC-induced effects such as hypothermia (Malone and Taylor, 2001), catalepsy (Egashira et al., 2007), anxiety, sociability and memory (Viñals et al., 2015). Therefore, we explored the role of serotonergic signalling on the amnesic-like effects of repeated low-dose THC 1. Of special interest were previous data showing that mice lacking the 5-HT2AR, or following pre-treatment with the 5-HT2AR antagonist MDL 100,907, prevented the acute effects of a high dose of THC in object recognition memory (Viñals et al., 2015). Using a similar pharmacological approach, we observed that the memory impairment produced by sub-chronic THC 1 mg/kg administration was totally prevented by 5-HT2AR antagonism by MDL. In addition, we described for the first time the involvement of 5-HT2AR in emotional memory impairment produced by THC since MDL blockade also prevented amnesic-like effects of THC in fear conditioning.

Based on these data, we hypothesized that CB1R/5-HT2AR heterodimers could underlie this sustained effect of THC. To test this we measured CB1R/5-HT2AR heterodimers in CA3, a hippocampal brain region in which this heterodimers have been previously described in mice (Viñals et al., 2015), and an area directly connected to CA1 region. We found a significant enhancement of CB1R/5-HT2AR heterodimers after sub-chronic THC 1 exposure and this modification was prevented by MDL 100,907 blockade of 5-HT2ARs. This result agrees with previous research in humans where enhanced expression of CB1R/5-HT2AR heterodimers was found in olfactory neuroepithelium cells of cannabis users (Galindo et al., 2018). Additionally, altered signalling in CB1R/5-HT2AR heterodimers also in olfactory neuroepithelium cells was reported in schizophrenia patients (Guinart et al., 2020) were a sensitized serotonergic responses are characteristic. Interestingly, when the entire population was considered, we found that a high level of CB1R/5HT2AR heterodimer expression in CA3 was associated with worse memory performance. Specifically, negative correlations were observed between the percentage of heterodimers and discrimation index in NOR task. This finding in also in agreement with a previous study when a significant negative correlation was stabilised between expression of CB1R/5-HT2AR heterodimers was found in olfactory neuroepithelium cells of cannabis users and their performance in cognitive tasks (Galindo et al., 2018). However, further studies addressing the specific relevance of the CB1R/5-HT2AR heterodimers will be important in understanding the cognitive deficits associated with repeated THC exposure, as well as the potential therapeutic potential of 5-HT2AR antagonists in preventing these deficits.

Together, our study indicates that even a low non-amnesic doses of THC can produce deleterious effects on memory function when administered repeatedly, and that these effects are accompanied by changes at the molecular and cellular levels involving the serotonergic system. Our data also implicate 5-HT2AR heterodimers in the effects of repeated low-dose THC on memory function.

## Acknowledgements

We are grateful to Marta Linares, Dulce Real, Raquel Martín, Jolita Jancyte and Francisco Porrón for expert technical assistance and the Laboratori de Neurofarmacologia-NeuroPhar for helpful discussion.

---

**Supplementary Figure 1.**
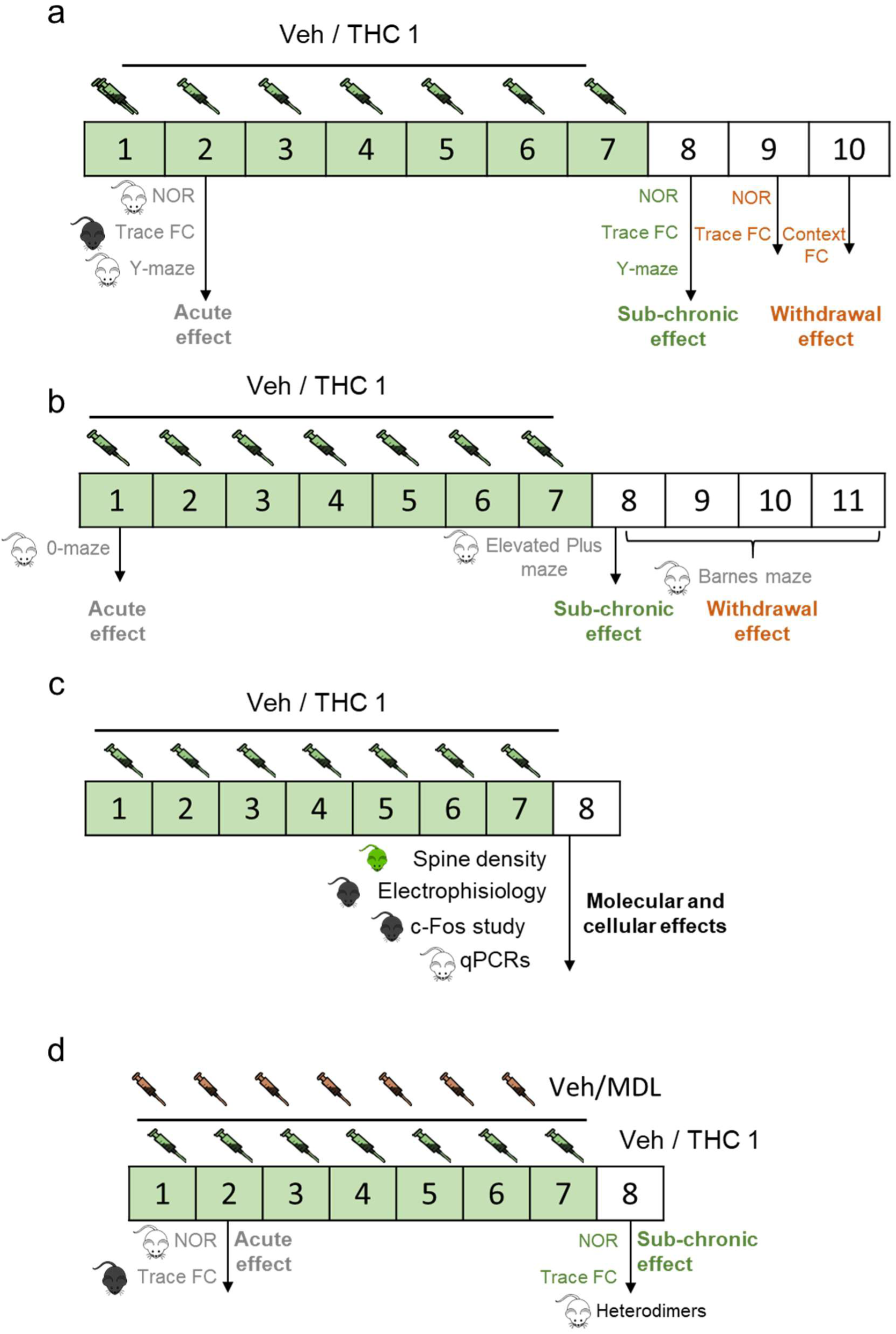
Timeline scheme of the behavioral tests, the treatments and animals used in the study. (a) Memory test performed to compare acute with sub-chronic effects of THC 1 were performed in the same way: the acute effect is measured 24h after the first administration and the sub-chronic 24h after the last administration, coinciding with the test phase of NOR task, Trace FC and Y-maze. In the case of treatment withdrawal effects, NOR and Trace FC test were performed 48h after the last administration and Context FC test 72h after last administration. (b) Additional behavioural task was used: anxiety-like behaviour was assessed in 0-maze 30 min after the first administration and in elevated plus maze 24h after the last administration. Spatial learning was measured using Barnes maze during 4 consecutive days starting 24h after the last sub-chronic administration. (c) For molecular and cellular effects studied samples were extracted 24h after the last administration. (d) For the evaluation of 5-HT2AR blockade with MDL, the pre-treatment was administered 20 min prior the THC 1 treatment and acute and sub-chronic affects were studied in NOR and Trace FC in the same way as previously.

**Supplementary Figure 2.**
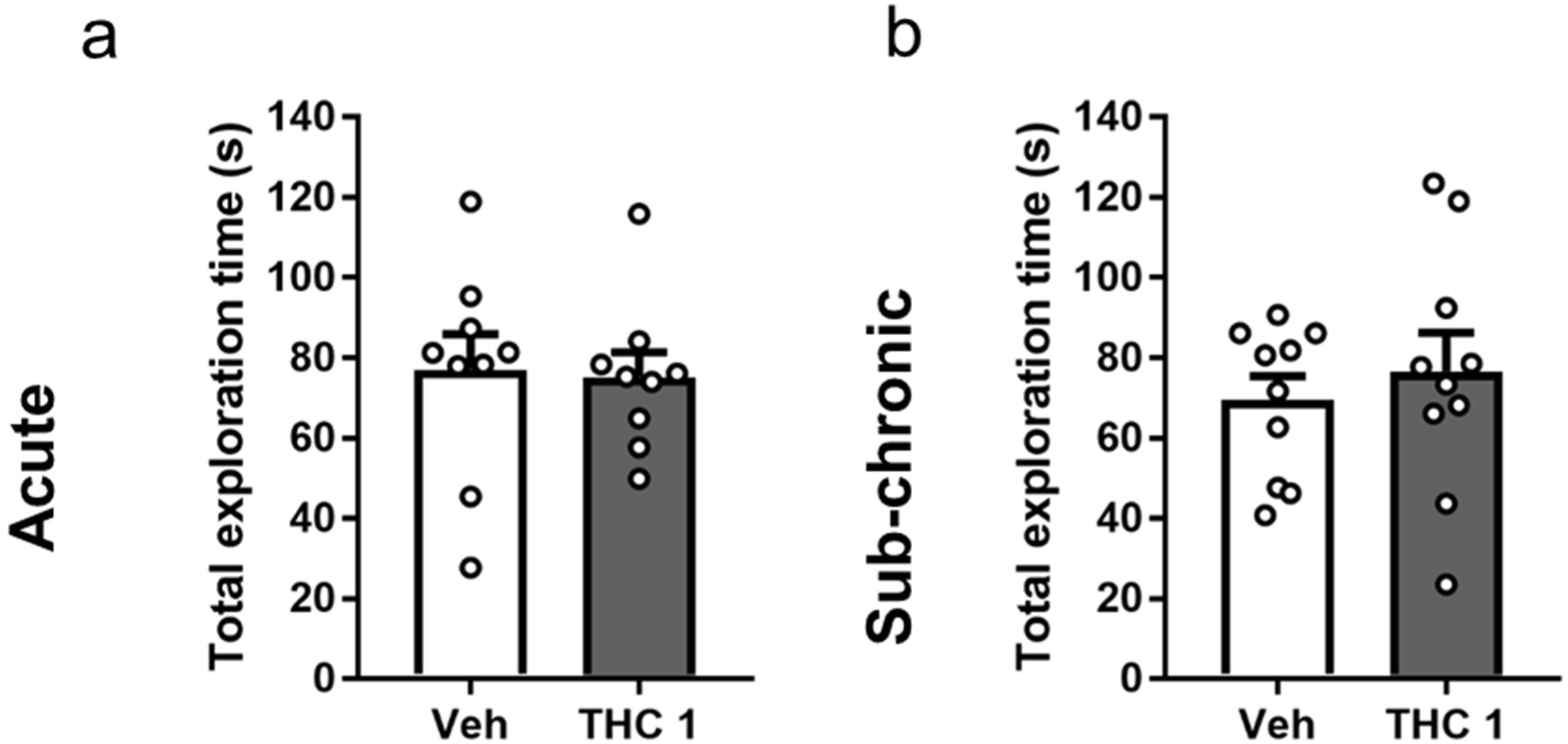
Total exploration time measurements in acute and sub-chronic NOR test. After administration of THC 1 no significant differences were observed neither in (a) acute (Veh n=9, THC 1 n=9) or (b) after the sub-chronic treatment (Veh n=10, THC 1 n=10) in total exploration times in NOR task. Statistical significance was calculated by Student’s t-test. Data are expressed as mean ± S.E.M.

**Supplementary Figure 3.**
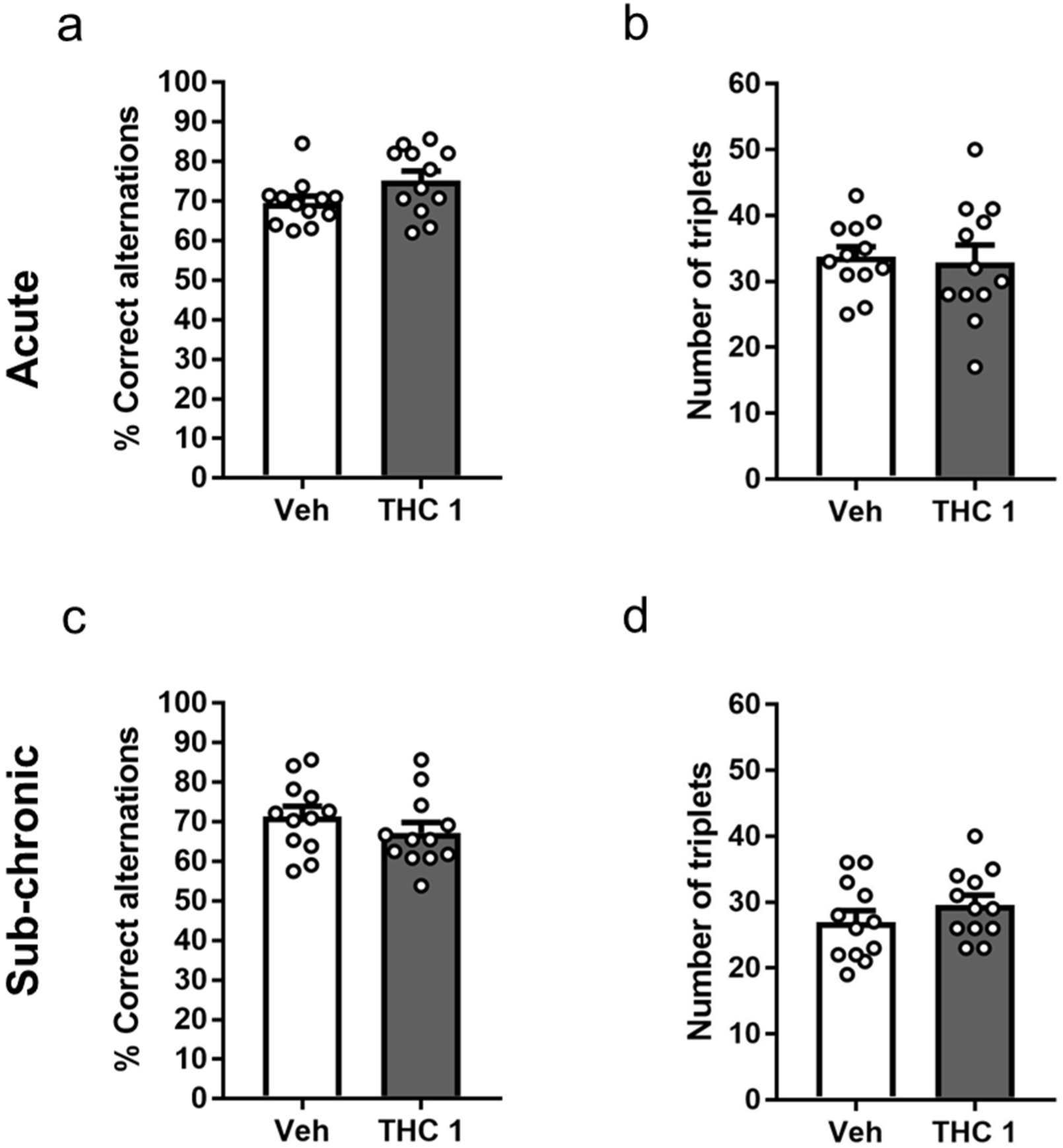
No effects of sub-chronic treatment with THC 1 in working-memory. After acute administration of THC 1 no significant differences were observed neither in (a) working-memory assessment Y-maze (Veh n=12, THC 1 n=12) or (b) number of triplets in Y-maze (Veh n=12, THC 1 n=12). After the sub-chronic treatment, no significant differences were observed either in (c) % of correct alternations (Veh n=12, THC 1 n=12) or (d) locomotion in Y-maze (Veh n=12, THC 1 n=12). Statistical significance was calculated by Student’s t-test. Data are expressed as mean ± S.E.M.

**Supplementary Figure 4.**
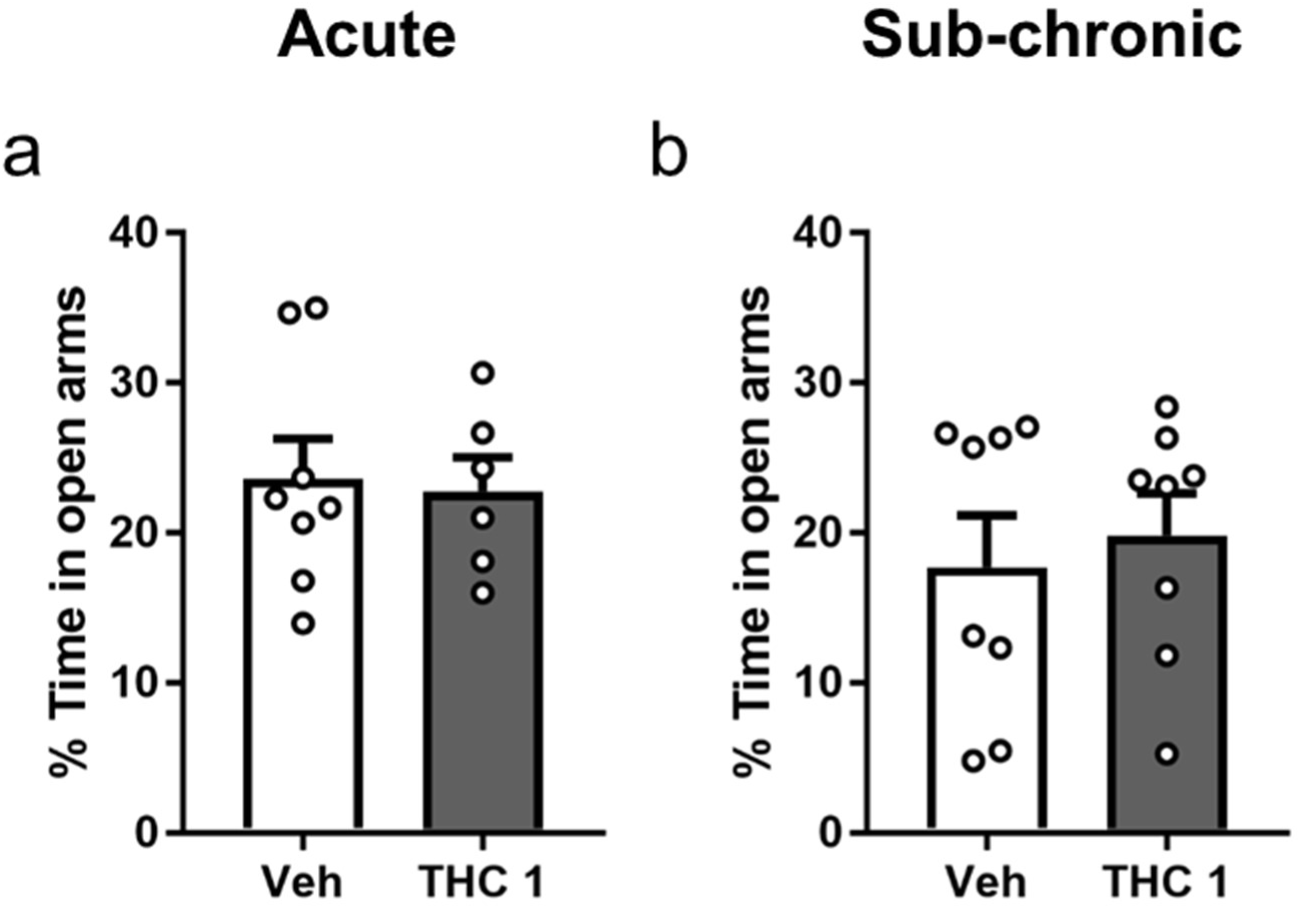
Exposure to THC 1 mg/kg do not modify anxiety-like behaviour. (a) 30 min after an acute injection of THC 1 mg/kg no differences in anxiety-like behaviour are observed in 0-maze (Veh n=8, THC 1 n=6). (b) After sub-chronic treatment with THC 1 mg/kg no effects are observed neither in anxiety-like behaviour in elevated plus maze (Veh n=8, THC 1 n=8). Statistical significance was calculated by Student’s t-test. Data are expressed as mean ± S.E.M.

**Supplementary Figure 5.**
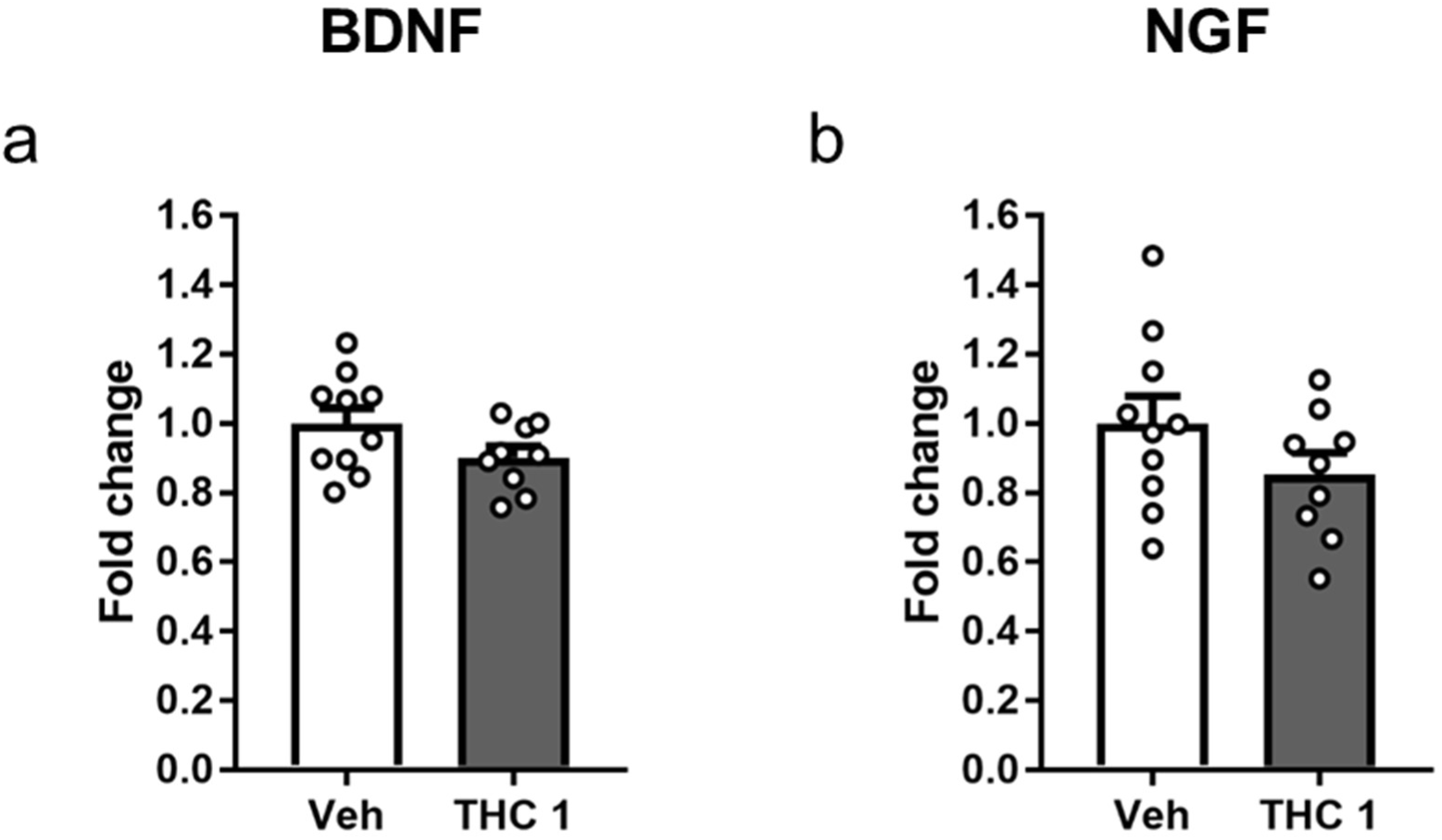
Neurotrophic factors after sub-chronic THC 1 exposure showed no changes at RNA level. qPCRs of the two principal neurotrophic factors in the hippocampus of vehicle and THC 1 animals revealed no significant differences neither in (a) BDNF (Veh n=10, THC 1 n=9) or (b) NGF (Veh n=10, THC 1 n=9). Statistical significance was calculated by Student’s t-test. Data are expressed as mean ± S.E.M.

**Supplementary Figure 6.**
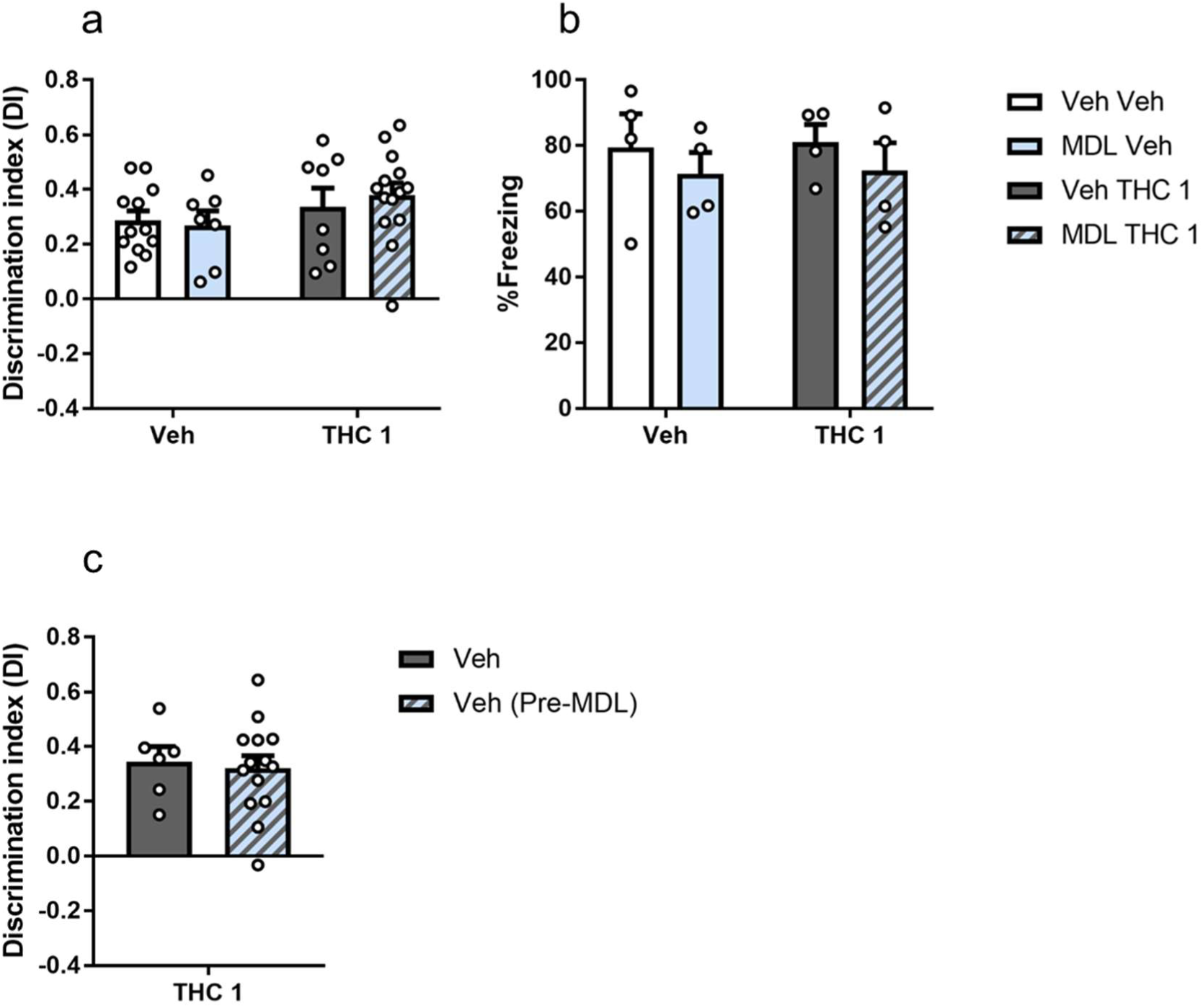
After an acute administration of THC 1 mg/kg or MDL 0.01 mg/kg no effects on memory are present. (a) Any of the treatments affect memory performance in an acute administration in (a) NOR task (Veh Veh n=12, MDL Veh n=7, Veh THC 1 n=8, MDL THC 1 n=14) and in (b) Trace fear conditioning (Veh Veh n=4, MDL Veh n=4, Veh THC 1 n=4, MDL THC 1 n=4). Statistical significance was calculated by Bonferroni post hoc test following two-way ANOVA. (c) No differences are present at memory performance at acute level in the NOR task in the groups in which the acute pre-treatment with MDL was tested. (Veh THC 1 n=6, Veh (Pre-MDL) THC 1 n=14). Statistical significance was calculated by Student’s t-test. Data are expressed as mean ± S.E.M.

**Supplementary Figure 7.**
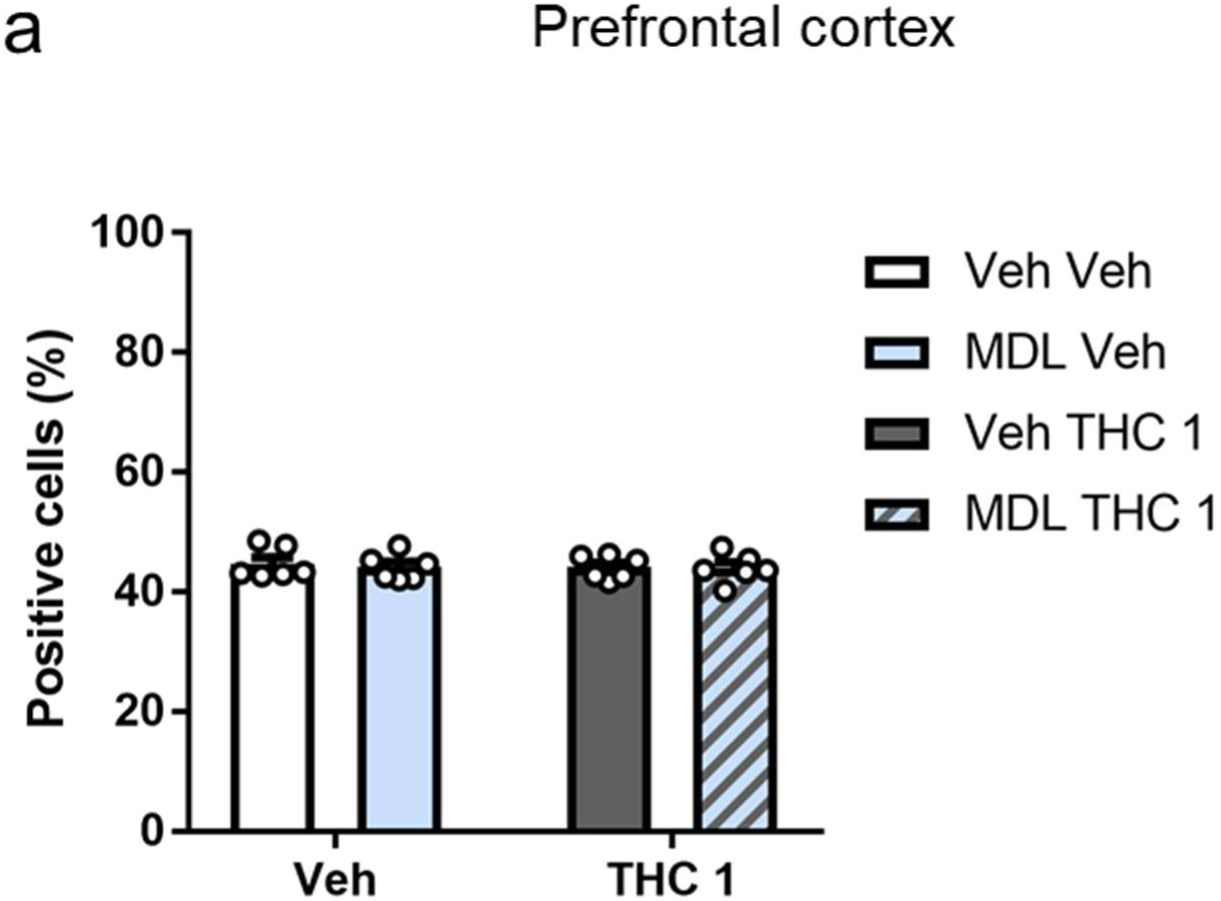
Heterodimers formed between CB1 and 5-HT2A receptor are not altered after THC 1 exposure in prefrontal cortex. (a) In prefrontal cortex no differences are observed in heterodimers levels between groups (Veh Veh n=6, MDL Veh n=6, Veh THC 1 n=6, MDL THC 1 n=6). Statistical significance was calculated by Bonferroni post hoc test following two-way ANOVA. Data are expressed as mean ± S.E.M.

